# Hippocampal–midcingulate connectivity is associated with representational integration during cognitive map updating

**DOI:** 10.64898/2026.07.16.738868

**Authors:** Jing Fu, Chengmei Huang, Ruimin Wang, Isao Hasegawa, Koji Jimura, Kiyoshi Nakahara

## Abstract

Cognitive maps organize experience into structured internal models that support goal-directed behavior. Although map formation has been extensively studied in the human hippocampus, the neural mechanisms supporting flexible updating of established maps remain unclear. We used fMRI to track the formation and updating of a spatial cognitive map across three days. Participants learned object–location associations and later updated them by remapping objects among fixed locations. Representational similarity analysis revealed two concurrent signatures in the right hippocampus: prior relational structure remained differentiated, whereas updated associations became integrated. Map updating engaged a frontoparietal network and modulated hippocampal–cortical connectivity. Critically, hippocampal–midcingulate connectivity was associated with the integration of updated associations, but not with the differentiation of prior relational structure. These results suggest that cognitive map updating involves coordinated hippocampal–cortical interactions that incorporate new information while preserving prior relational structure, reflecting a balance of stability and plasticity in internal models.

## Introduction

The brain organizes experience into structured internal models, known as cognitive maps, which provide a foundational scaffold for planning and goal-directed behavior^1–3^. Through spatial navigation, humans acquire not only the discrete locations of individual landmarks^4–6^, but also the relational knowledge linking them^7,8^, thereby constructing a holistic representation of the spatial environment^9^. However, real-world environments are rarely static, presenting a fundamental challenge: how these established cognitive maps are flexibly updated and reorganized when previously learned spatial relationships change.

Central to constructing such relational structures is the hippocampus, which has long been implicated in the formation and retrieval of cognitive maps and episodic memories^10–13^. One hypothesized hippocampal mechanism is pattern separation, which transforms highly similar input patterns into more distinct, non-overlapping representations^14,15^, thereby mitigating interference among events with shared features^16^. In parallel, the hippocampus also supports relational integration, binding information across separate events that share common elements to form structured representations^2,17,18^. Recent work demonstrates that hippocampal mechanisms, such as pattern separation and integration, jointly support the emergence of structured knowledge during map formation^19^.

Once a cognitive map is established, however, updating that map is not equivalent to learning about a new environment from scratch. Instead, it requires integrating new information—sometimes conflicting with prior knowledge—into existing representations by modifying relational structures^20^. Existing representations can interfere with the integration of new information, whereas indiscriminate updating can render prior knowledge vulnerable to erasure^21–23^. Successful updating therefore requires resolving the tension between preserving prior knowledge and incorporating new experience, a challenge formalized as the stability–plasticity dilemma^22,24^. Recent computational models propose that persistent "traces of experience" within hippocampal output structures facilitate cognitive map updating through mismatch-dependent integration of new spatial contingencies^25^. Despite these insights, it remains unclear how hippocampal representations accommodate new, conflicting information while preserving prior relational structure during map updating.

Beyond intrinsic hippocampal mechanisms, successful cognitive map updating is likely to require coordinated interactions across distributed hippocampal–cortical networks. The parietal cortex has been associated with the encoding and transformation of spatial representations across reference frames^26^, with rodent work further highlighting a hippocampal–parietal network involved in coordinating stored spatial knowledge with prospective, goal-directed representations^27^. Moreover, hippocampal–medial prefrontal circuits in humans have been particularly implicated in flexible memory updating, a process by which established memories are modified to incorporate novel content^28^. More dorsal medial frontal regions, including the midcingulate cortex, have been implicated in cognitive control^29,30^—functions that may be recruited when incoming relational information conflicts with an established representational structure. Nevertheless, these lines of work largely characterize hippocampal–cortical interactions during novel encoding, spatial transformation, or general memory modification. How distributed hippocampal–cortical systems support the updating of established cognitive maps remains poorly characterized.

To address these gaps, we employed a three-day fMRI paradigm to track the neural mechanisms underlying the formation and updating of spatial cognitive maps (Fig. 1a). Inspired by the Objects in Updated Locations paradigm^31–33^, we developed a Spatial Map Updating Task in which objects were anchored to a set of fixed spatial nodes. Notably, by remapping objects among these fixed nodes, participants rearranged existing object–location associations within a stable spatial scaffold rather than encoding entirely novel coordinates, thereby isolating adaptive relational restructuring from *de novo* map formation. Unlike standard virtual navigation paradigms that rely on continuous path integration in large-scale environments^8,34^, this approach reduces confounds associated with complex sensory cues and strategy use. Participants learned object–location associations on Day 1 to establish an initial spatial map and subsequently updated the map on Day 2 according to predefined shift cues. To track representational changes across map formation and updating, neural responses were measured during an incidental picture viewing task (PVT) at three time points—prior to learning (Day 1), after learning but before updating (Day 2), and after updating (Day 3).

**Fig. 1:**
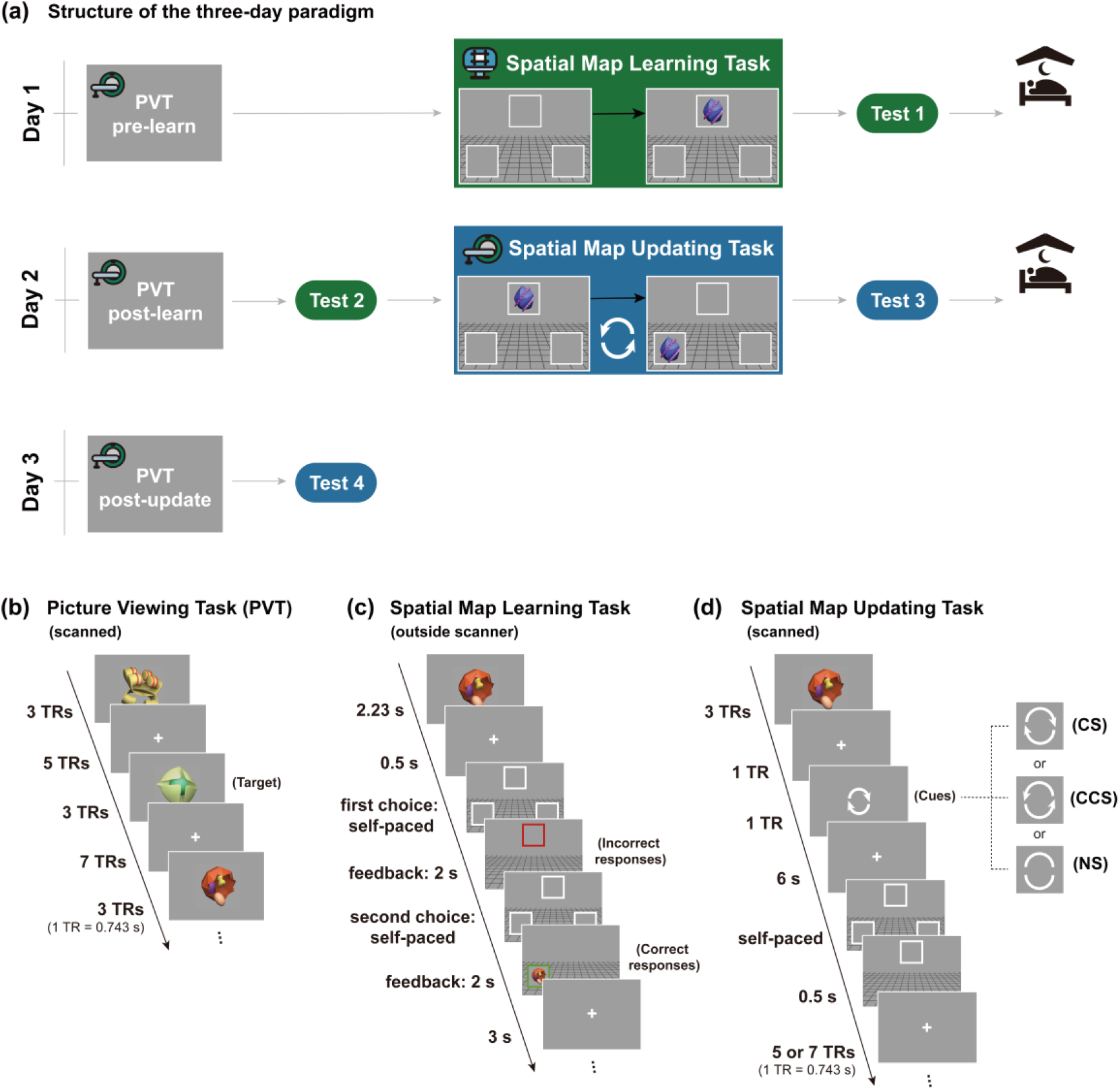
Experimental overview and task design. (a) Structure of the three-day paradigm. On Day 1, participants completed a baseline picture viewing task scan (PVT pre-learn), followed by a behavioral spatial map learning task and a post-learning memory test (Test 1). On Day 2, participants completed a post-learning PVT scan (PVT post-learn), a post-learning retention test (Test 2), an fMRI spatial map updating task, and a post-updating memory test (Test 3). On Day 3, participants completed a final post-updating PVT scan (PVT post-update) and a post-updating retention test (Test 4). (b) Picture Viewing Task (PVT). Nine task-relevant objects and one target object were presented in a pseudorandomized sequence that was identical across all three sessions. Participants made a button press only when the target object appeared. (c) Spatial Map Learning Task. Participants acquired object–location associations through trial-and-error learning with immediate deterministic feedback. Correct responses were indicated by a green frame and presentation of the object, whereas incorrect responses were indicated by a red frame without object presentation. Following an incorrect choice, participants continued selecting locations until the correct location was identified. (d) Spatial Map Updating Task. Participants updated the previously learned object–location associations according to one of three cues: clockwise shift (CS), counterclockwise shift (CCS), or non-shift (NS). Each object was paired with a single cue throughout the task. The CS and CCS cues instructed participants to move the object’s associated location by one position in the corresponding direction, whereas the NS cue indicated no change. No feedback was provided; instead, the selected frame remained visible for 0.5 s to acknowledge the response.

Using this paradigm, we investigated how hippocampal representations are reorganized during cognitive map updating and whether hippocampal–cortical functional connectivity is associated with this representational reorganization. We found evidence for concurrent integration of the updated map and differentiation of the original map within the right hippocampus. Map updating further engaged a distributed frontoparietal network, within which hippocampal–midcingulate connectivity was selectively associated with the integration of updated associations rather than the differentiation of prior relational structure. Together, these findings suggest a hippocampal–cortical account of flexible cognitive map updating in which new information is incorporated while prior relational structure is retained.

## Results

We employed a three-day experimental paradigm (Fig. 1a) to investigate cognitive map formation and subsequent updating.

On Day 1, participants first underwent a baseline PVT during fMRI scanning, in which they viewed ten novel objects and responded to a designated target object. The target object was exclusive to the PVT, whereas the remaining nine objects were used in subsequent learning tasks. Participants then completed a spatial map learning task outside the scanner, in which the nine task-relevant objects were assigned to three locations, with three objects associated with each location. After learning these object–location associations through repeated training trials, participants completed an immediate memory test to verify acquisition of the initial associations.

On Day 2, participants underwent a second PVT scan and then completed a memory test outside the scanner to assess retention of the object–location associations acquired on Day 1. Participants then returned to the scanner to complete a spatial map updating task. In this task, each object was consistently paired with one of three cue types—clockwise shift (CS), counterclockwise shift (CCS), or non-shift (NS; Supplementary Fig. 1b)—such that each object was always subject to the same type of manipulation. CS and CCS trials required participants to update the previously learned object–location associations and were therefore grouped as shift trials, corresponding to the updating condition. NS trials involved no change in associations and served as a within-task control condition. Following the updating task, participants completed an immediate post-update memory test outside the scanner to assess acquisition of the updated associations.

On Day 3, participants underwent a final PVT scan and then completed a memory test outside the scanner to assess retention of the updated associations acquired on Day 2.

### Behavioral performance

#### Picture Viewing Task (fMRI session)

Nine task-relevant objects and one target object were each presented 12 times in a pseudorandomized order (Fig. 1b). The presentation sequence was identical across the pre-learning (PVT pre-learn; Day 1), post-learning (PVT post-learn; Day 2), and post-updating (PVT post-update; Day 3) sessions. Participants responded via button press whenever the target object appeared. Analyses included the 43 participants who completed all three PVT sessions (Table 1). Accuracy was near ceiling across sessions (PVT pre-learn [mean ± SD]: 99.1% ± 2.4%; PVT post-learn: 99.8% ± 0.7%; PVT post-update: 99.8% ± 0.7%), indicating sustained attention to the task.

**Table 1:** Initial and final sample sizes for each task.

| Task / Analysis | Initial N | Excluded N | Exclusion Criteria | Final N |
| --- | --- | --- | --- | --- |
| PVT (Day 1) | 45 | 0 | — | 45 |
| Spatial Map Learning Task | 45 | 0 | — | 45 |
| PVT (Day 2) | 45 | 0 | — | 45 |
| Spatial Map Updating Task | 45 | 6 | Withdrawal ($n = 1$);<br>Excessive motion ($n = 5$) | 39 |
| PVT (Day 3) | 45 | 2 | Absence | 43 |
| Connectivity–Representation<br>Analysis | N/A | — | 39 (Updating Task) ∩ 43 (PVT Day 3) | 38 |

*Note.* Of the two participants who did not complete PVT (Day 3), one had previously withdrawn from the study because of physical discomfort during the spatial map updating session, and the other was absent on Day 3. The connectivity–representation analysis (final N = 38) included participants with valid data from both the spatial map updating task and PVT (Day 3).

#### Spatial Map Learning Task (behavioral session)

We examined response accuracy during the spatial map learning task on Day 1 to assess initial learning performance (outside the scanner; Fig. 1c). The nine objects were presented in mini-blocks such that all stimuli appeared once before any repetition occurred. Across five learning runs (10 mini-blocks in total), participants (*n* = 45; Table 1) showed a progressive increase in response accuracy. Performance exceeded chance level (33.3%) in the first mini-block (mini-block 1 [mean ± SD]: 40.5% ± 16.3%; *t*(44) = 3.09, *p* = 0.003, *d* = 0.46, one-sample *t*-test) and reached near-ceiling levels by the final mini-block (mini-block 10: 99.3% ± 3.7%; *t*(44) = 121.12, *p* < 0.001, *d* = 18.06). Participants subsequently completed an immediate memory test consisting of four mini-blocks and demonstrated high retrieval accuracy (99.3% ± 4.2%), indicating successful acquisition of the object–location associations.

To assess overnight retention of the learned associations, participants completed a retention test on Day 2, administered after the picture viewing task to avoid priming the memory representations subsequently probed during scanning. Retention accuracy on Day 2 (98.8% ± 4.0%) did not significantly differ from immediate memory test performance on Day 1 (99.3% ± 4.2%; *t*(44) = 1.39, *p* = 0.173, *d* = 0.21, paired-samples *t*-test).

#### Spatial Map Updating Task (fMRI session)

Of the 45 participants, 39 completed the spatial map updating task during fMRI scanning on Day 2 (Fig. 1d and Table 1). In this analytic sample, updating performance was highly accurate for both shift trials (97.2% ± 5.7%) and non-shift trials (97.0% ± 6.6%), with no significant difference between conditions (*t*(38) = 0.24, *p* = 0.814, *d* = 0.04, paired-samples *t*-test). Reaction times for correct trials were also comparable between conditions (shift: 583 ± 162 ms; non-shift: 609 ± 210 ms; *t*(38) = −1.74, *p* = 0.091, *d* = −0.28, paired-samples *t*-test). An immediate post-update memory test further confirmed high retrieval accuracy (94.6% ± 13.5%). These results indicate that participants successfully implemented the updating rules and acquired the updated associations.

A delayed memory test administered on Day 3 showed high retrieval accuracy (91.5% ± 17.4%) but revealed a significant decline relative to the immediate post-update test on Day 2 (94.6% ± 13.5%; *t*(38) = 2.67, *p* = 0.011, *d* = 0.43, paired-samples *t*-test). Nevertheless, performance remained high, indicating that the updated associations were largely retained across days.

### Hippocampal representational changes following learning and updating

To examine how hippocampal representations changed across spatial learning and subsequent updating, we extracted voxel-level activity patterns from the left and right hippocampus and quantified representational change using differences in representational dissimilarity matrices (ΔRDMs). ΔRDMlearn, calculated as the post-learning PVT RDM minus the pre-learning PVT RDM, captured representational changes associated with initial learning. ΔRDMupdate, calculated as the post-updating PVT RDM minus the pre-learning PVT RDM, captured cumulative representational changes following updating (Fig. 2a; see Methods).

**Fig. 2:**
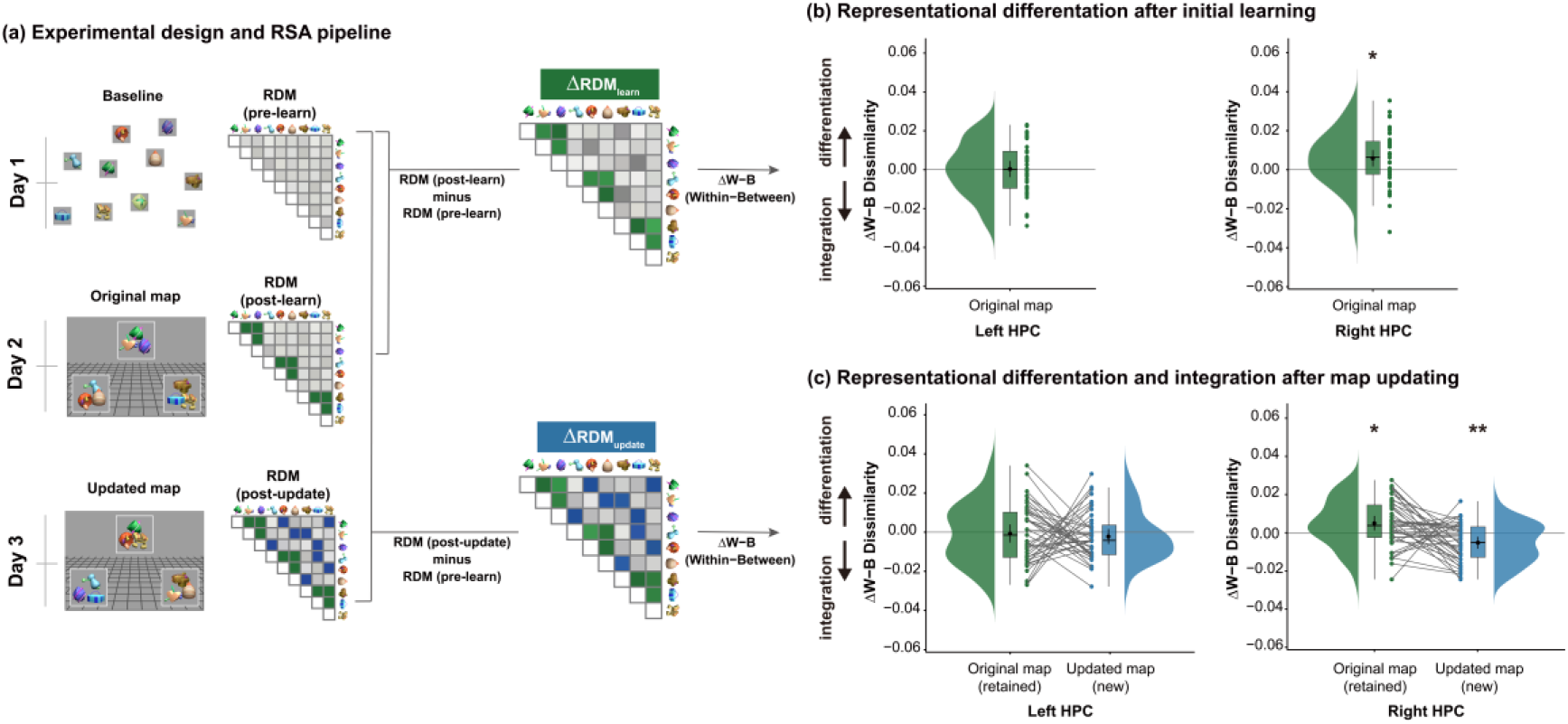
Hippocampal representational reorganization across learning and updating. (a) Experimental design and RSA pipeline. Voxel-level activity patterns within the left and right hippocampus (HPC) were extracted to construct representational dissimilarity matrices (RDMs) for each phase (pre-learning, post-learning, and post-updating). Change RDMs (ΔRDM) were computed by subtracting the pre-learning RDM from each subsequent phase. A within–between dissimilarity index (ΔW−B) was derived from each ΔRDM by contrasting dissimilarity changes for within-location versus between-location object pairs. In these ΔRDM, green cells denote object pairs sharing an original map location, blue cells denote object pairs sharing an updated map location, and gray cells correspond to between-location object pairs. (b) Representational differentiation after initial learning. Raincloud plots show ΔW−B for the original map structure (ΔRDMlearn: RDM post-learn minus RDM pre-learn) in the left and right hippocampus. Positive ΔW−B values indicate greater increases in dissimilarity for within-location than between-location object pairs. The right hippocampus showed significant representational differentiation, whereas no significant effect was observed in the left hippocampus. (c) Representational differentiation and integration after map updating. Raincloud plots show ΔW−B for the original map structure retained from initial learning and the newly updated map structure (ΔRDMupdate: RDM post-update minus RDM pre-learn) in the left and right hippocampus. In the right hippocampus, the original map structure showed significant representational differentiation, whereas the updated map structure showed significant representational integration. No significant effects were observed in the left hippocampus. Linked lines connect each participant’s ΔW−B values for the retained original and updated map structures. Error bars denote mean ± 95% CI. \**p* < 0.05, \*\**p* < 0.01.

From each ΔRDMs, we derived a within–between dissimilarity change index (ΔW−B), defined as the mean change in dissimilarity for within-location object pairs minus that for between-location object pairs. Within-location pairs comprised objects occupying the same location, whereas between-location pairs comprised objects occupying different locations. Positive ΔW−B values indicate representational differentiation, reflecting a greater increase in dissimilarity for within-location than between-location pairs, whereas negative ΔW−B values indicate representational integration, reflecting a smaller increase (or relative decrease) in dissimilarity for within-location pairs than between-location pairs.

### Initial learning elicits representational differentiation

We examined representational changes following initial learning (ΔRDMlearn) using one-sample *t*-tests against zero (*n* = 43; Table 1). The right hippocampus exhibited a significant increase in ΔW−B (mean ± SD: 0.0059 ± 0.0137; *t*(42) = 2.80, *p* = 0.008, FDR-corrected *p* = 0.015, *d* = 0.43), indicating representational differentiation of the original map structure (Fig. 2b). No significant effect was observed in the left hippocampus (mean ± SD: 0.0003 ± 0.0136; *t*(42) = 0.16, *p* = 0.874, *d* = 0.02; Fig. 2b). A paired-samples *t*-test further confirmed that ΔW−B values were significantly greater in the right than in the left hippocampus (*t*(42) = 2.09, *p* = 0.043, *d* = 0.32).

### Updating elicits both representational differentiation and integration

We next quantified representational changes following updating (ΔRDMupdate). Because updating may give rise to coexisting representations of the original and updated relational structures, ΔW−B was computed separately for two sets of object pairs: one based on the original object–location mappings and the other based on the updated mappings. This allowed us to assess representational change with respect to both the original and updated map structures.

In the right hippocampus, ΔW−B for the original map was significantly above zero (mean ± SD: 0.0050 ± 0.0124; *t*(42) = 2.64, *p* = 0.012, FDR-corrected *p* = 0.023, *d* = 0.40), indicating that the original map structure remained differentiated following updating (Fig. 2c, green box). In contrast, ΔW−B for the updated map was significantly below zero (mean ± SD: −0.0053 ± 0.0103; *t*(42) = −3.36, *p* = 0.002, FDR-corrected *p* = 0.003, *d* = −0.51), indicating representational integration, with relatively greater similarity among object pairs sharing the same updated location (Fig. 2c, blue box). Together, these results suggest that original and updated relational structures coexisted within right hippocampal representations.

No significant differentiation or integration effects were observed in the left hippocampus (Supplementary Table 1). Although these effects were numerically stronger in the right hippocampus, inter-hemispheric comparisons did not reach significance for either effect (differentiation: *t*(42) = 1.74, *p* = 0.089, *d* = 0.27; integration: *t*(42) = −1.26, *p* = 0.214, *d* = −0.19, paired-samples *t*-tests).

### Model-based RSA provides complementary evidence for original and updated map structures in right hippocampal representations

Complementing the within–between dissimilarity analysis described above, we constructed model RDMs capturing the relational structure of the original and updated spatial maps and computed model fits as Spearman correlations between model RDMs and the neural ΔRDMs derived from crossnobis distances (ΔRDMlearn and ΔRDMupdate; see Methods). The neural ΔRDMs represented changes in pairwise dissimilarity, whereas the model RDMs coded within-location pairs as 1 and between-location pairs as 0. A positive model–neural correlation therefore indicated a relative increase in dissimilarity among within-location compared with between-location object pairs, consistent with representational differentiation.

Conversely, a negative correlation indicated a relative decrease in dissimilarity among within-location compared with between-location object pairs, consistent with representational integration.

Following initial learning, right hippocampal ΔRDMs showed a significant positive fit with the original model RDM (mean Fisher-*z* fit = 0.068, *t*(42) = 3.26, *p* = 0.002, FDR-corrected *p* = 0.004, *d* = 0.50), indicating that learning-related representational change was aligned with differentiation according to the original map structure. Following updating, right hippocampal ΔRDMs showed a significant positive fit with the original model RDM (mean Fisher-*z* fit = 0.054, *t*(42) = 3.07, *p* = 0.004, FDR-corrected *p* = 0.007, *d* = 0.47), indicating persistent differentiation according to the original map structure. In contrast, right hippocampal ΔRDMs showed a significant negative fit with the updated model RDM (mean Fisher-*z* fit = −0.052, *t*(42) = −3.09, *p* = 0.003, FDR-corrected *p* = 0.007, *d* = −0.47), indicating integration according to the updated map structure. No significant corresponding effects were observed in the left hippocampus (Supplementary Table 5).

To assess whether each updating-related effect reflected variance uniquely associated with its respective map structure, we repeated the analysis using partial Spearman correlations while controlling for the other model RDM (see Methods). The two model RDMs were modestly negatively correlated (Spearman’s ρ = −0.33). Both associations were attenuated and did not survive FDR correction (original model: *t*(42) = 2.22, *p* = 0.032, FDR-corrected *p* = 0.063, *d* = 0.34; updated model: *t*(42) = −2.16, *p* = 0.037, FDR-corrected *p* = 0.074, *d* = −0.33), indicating that representational changes associated with the original and updated map structures were not fully independent (Supplementary Table 6).

Taken together, these model-based RSA results provide complementary evidence supporting the ΔW−B findings. Specifically, right hippocampal representational change reflected differentiation according to the original map structure during initial learning. After updating, this differentiation according to the original structure persisted, and was accompanied by concurrent integration according to the updated map structure.

### Whole-brain activation patterns during the spatial map updating task

We performed a whole-brain univariate general linear model (GLM) analysis contrasting shift trials (CS and CCS) with non-shift trials (NS; voxel-wise FDR-corrected *p* < 0.01; see Methods) to identify brain regions associated with spatial map updating. Analyses included 39 participants with valid fMRI data after head-motion exclusions (Table 1). Because shift and non-shift trials showed comparable accuracy and correct-trial reaction times, the shift versus non-shift contrast was unlikely to be driven primarily by differences in task performance or time-on-task.

### Spatial map updating preferentially engages a distributed frontoparietal–cingulate–cerebellar network

Map updating (shift > non-shift) elicited widespread activations across frontal, parietal, occipital, and cerebellar regions (Fig. 3a). In the frontal regions, significant activations were observed in the bilateral precentral gyrus (PreCG), the superior frontal gyrus (SFG), the frontal eye fields (FEF), the supplementary motor area (SMA), the midcingulate cortex (MCC), the insular cortex (IC), and the left inferior frontal gyrus, pars triangularis (IFGtri).

**Fig. 3:**
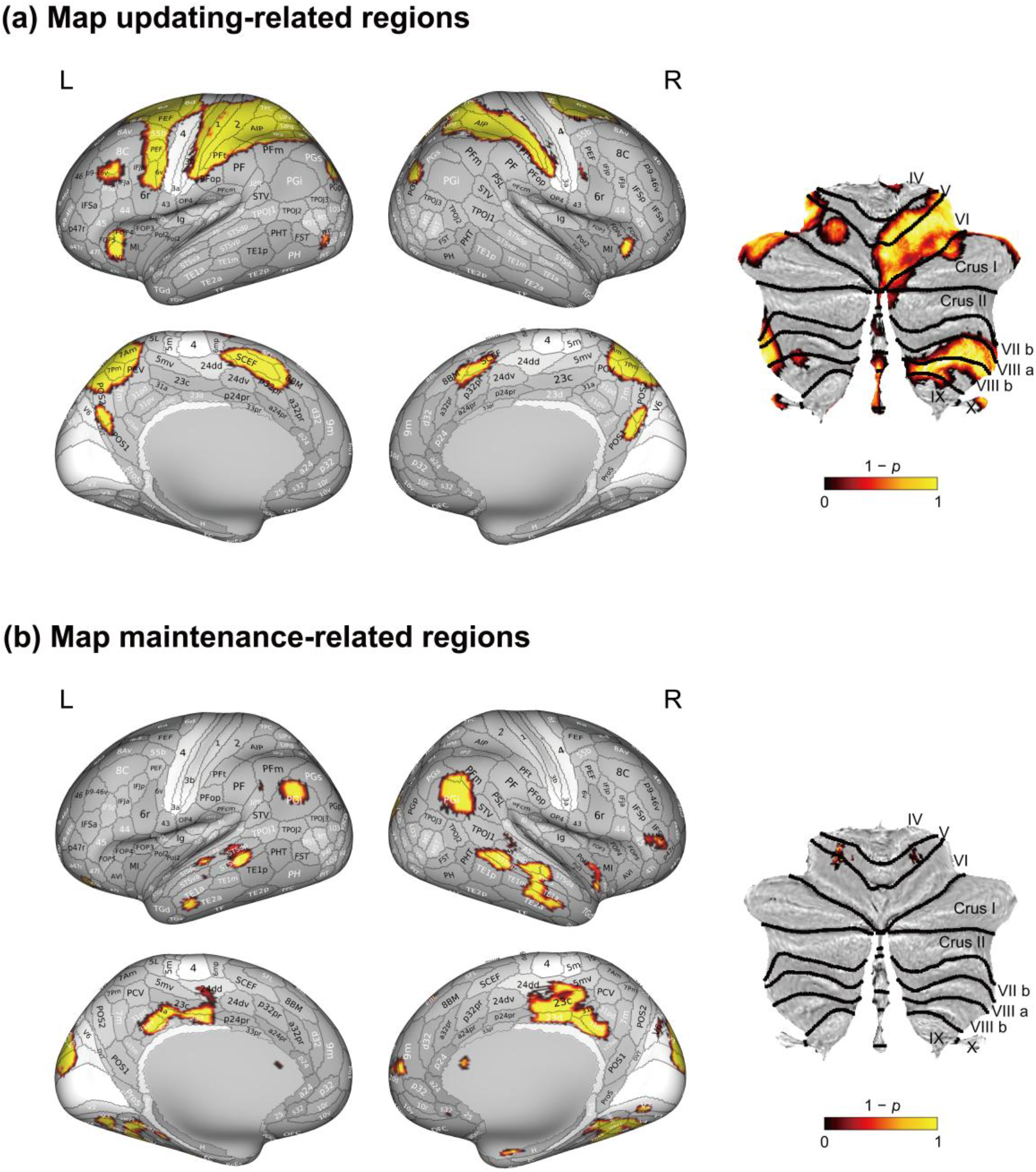
Brain activations associated with cognitive map updating and maintenance. (a) Map updating-related regions (shift > non-shift). Widespread activation was observed across frontal regions (PreCG, SFG, FEF, SMA, MCC, IC, IFGtri), parietal regions (SPL/IPL, IPS, precuneus), MOG, and cerebellum. The cerebellar flatmap (right) shows activations centered on bilateral lobule VI and Crus I, extending into lobule VIII, lobule X, and vermis. (b) Map maintenance-related regions (non-shift > shift). Greater activation was observed in parietal regions (AG), temporal regions (MTG, ITG, FG), occipital regions (cuneus, SOG, lingual gyrus), MCC, right IFGoper, and right frontal pole. Lateral (top rows) and medial (bottom rows) surfaces of the inflated brains are displayed for both the left (L) and right (R) hemispheres, alongside flattened cerebellar representations in SUIT space. All activation maps are thresholded at a voxel-wise *p* < 0.01 (FDR corrected), derived from a non-parametric one-sample *t*-test with 10,000 permutations (color bars indicate 1 − *p* values). Abbreviations: PreCG, precentral gyrus; SFG, superior frontal gyrus; FEF, frontal eye fields; SMA, supplementary motor area; MCC, midcingulate cortex; IFGtri, inferior frontal gyrus, pars triangularis; IC, insular cortex; SPL, superior parietal lobule; IPL, inferior parietal lobule; IPS, intraparietal sulcus; AG, angular gyrus; MTG, middle temporal gyrus; ITG, inferior temporal gyrus; SOG, superior occipital gyrus; MOG, middle occipital gyrus; FG, fusiform gyrus; IFGoper, inferior frontal gyrus, pars opercularis.

Parietal activations included bilateral clusters in the superior and inferior parietal lobules (SPL/IPL), the intraparietal sulcus (IPS), and the precuneus (PCun). Additional clusters were identified in the bilateral middle occipital gyrus (MOG) and the left thalamus (Thal).

Cerebellar engagement was also observed, with prominent activations in bilateral lobule VI and Crus I, extending into posterior cerebellar regions including lobule VIII, lobule X, and vermis.

### Spatial map maintenance preferentially engages posterior occipitotemporal and angular gyrus regions

The non-shift > shift contrast revealed greater activation in parietal, temporal, and occipital cortices (Fig. 3b). Parietal activations were observed in the bilateral angular gyrus (AG), alongside temporal clusters in the bilateral middle temporal gyrus (MTG), the inferior temporal gyrus (ITG), and the fusiform gyrus (FG). Occipital regions included the bilateral cuneus (Cun), the superior occipital gyrus (SOG), and the lingual gyrus (LG). Additional clusters were identified in the bilateral MCC, the right inferior frontal gyrus, pars opercularis (IFGoper), and the right frontal pole (FPole).

Although MCC clusters were observed in both contrasts, their spatial distributions differed. Updating-related activation was located more anteriorly along the medial frontal cortex (MNI peak voxel coordinates: 14, 16, 36), in the region of the dorsal anterior MCC bordering the pre-supplementary motor area, whereas maintenance-related activation was centered in a more posterior MCC region (MNI peak voxel coordinates: −6, −20, 42; Supplementary Table 2).

### No differential activation across remapping directions

We contrasted CS and CCS conditions to examine whether the direction of spatial remapping engaged distinct neural systems. No suprathreshold clusters were detected, suggesting no reliable direction-specific differences in activation at the present threshold. Across all contrasts, no suprathreshold hippocampal clusters were identified in the whole-brain analyses.

### Task-modulated functional connectivity during the spatial map updating task

We performed a generalized psychophysiological interaction (gPPI) analysis to examine task-modulated functional connectivity between the hippocampus and brain regions engaged during map updating (see Methods). Thirty-two spherical ROIs were defined around peak activation coordinates identified in the group-level shift > non-shift contrast (Supplementary Table 2), collectively constituting the map-updating network. A two-tailed *t*-test with FDR correction (*p* < 0.05) identified 94 significant connections within this network (Fig. 4a; full list in Supplementary Table 3). Parietal ROIs exhibited the greatest number of significant connections, particularly with frontal ROIs (31 connections), other parietal ROIs (26 connections), and occipital ROIs (9 connections). Among hippocampal connections, only the right hippocampus showed significant task-modulated connectivity with the right midcingulate cortex (*β* = 0.209, *t*(38) = 2.83, *p* = 0.007, FDR-corrected *p* = 0.034). No significant connectivity was observed for the left hippocampus.

**Fig. 4:**
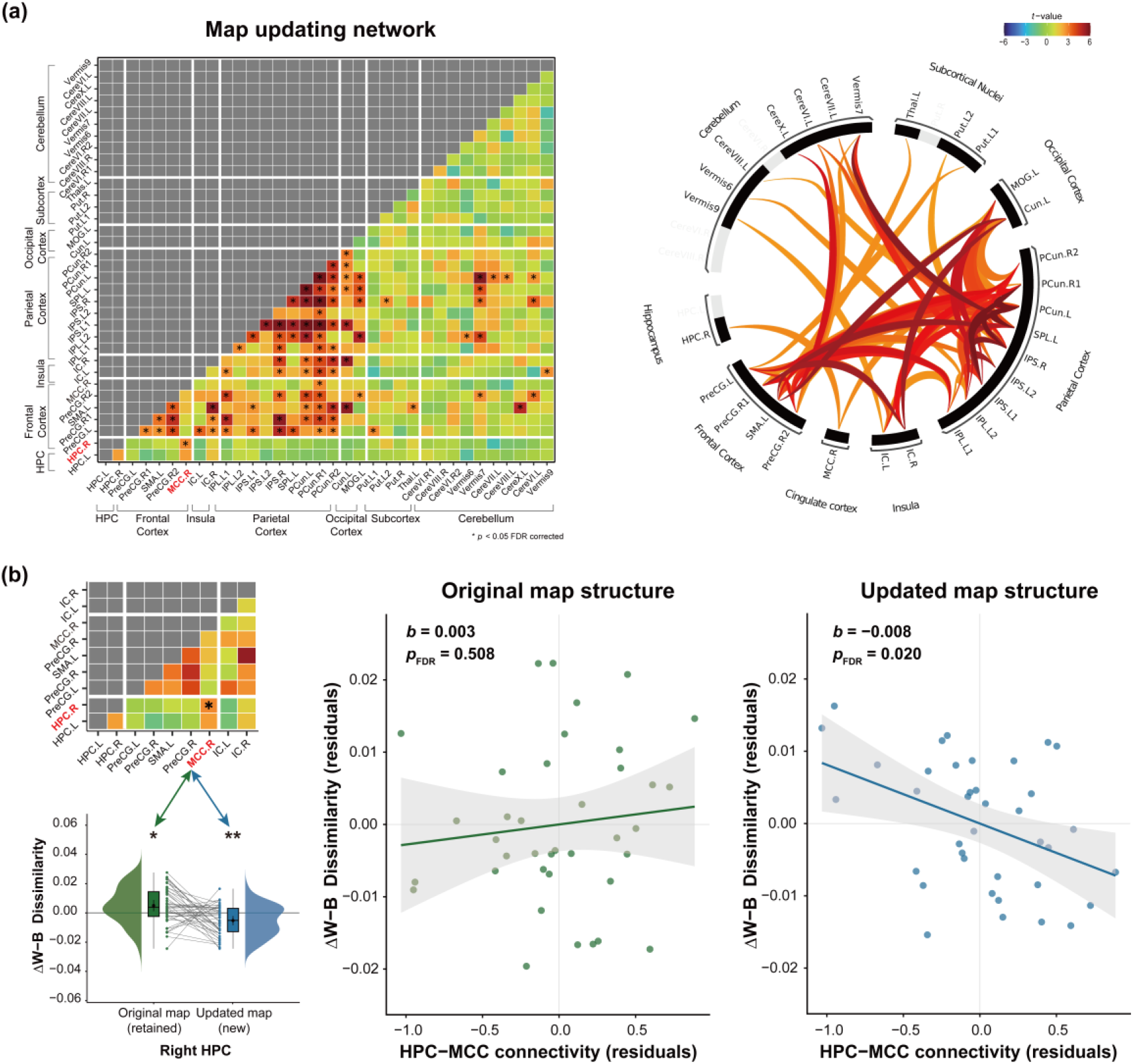
Task-based functional connectivity and its association with representational reorganization. (a) Map updating network and functional interactions. The heatmap (left) illustrates task-modulated functional connectivity among regions identified by the map-updating contrast (shift > non-shift), with color intensity reflecting *t*-values. Red labels on the axes indicate the right hippocampus (HPC.R) and right midcingulate cortex (MCC.R), highlighting the significant task-dependent connection (asterisk). The connectogram (right) is thresholded to highlight significant functional connectivity (*p* < 0.05, FDR corrected). (b) Hippocampal–MCC connectivity is associated with updated relational structure. Top left: Mini-connectivity matrix focusing on the hippocampal–cortical subnetwork. Bottom left: Raincloud plot showing the distribution of representational reorganization indices (ΔW−B) for the original map (retained) and the updated map (new), extracted from the right hippocampus. Middle and right: Partial regression scatter plots show the association between hippocampal–MCC functional connectivity (residuals) and representational reorganization indices ΔW−B (residuals) for the original map structure (middle; *b* = 0.003, *p*FDR= 0.508) and the updated map structure (right; *b* = −0.008, *p*FDR = 0.020), after controlling for mean FD. Shaded regions denote 95% confidence intervals. The *p* values were corrected across the original- and updated-map regression models using the Benjamini–Hochberg false discovery rate procedure.

In contrast, the map maintenance-related network, defined from the non-shift > shift contrast and comprising twenty-four ROIs (Supplementary Table 2), showed a comparatively sparse pattern of functional connectivity. Only three ROI-to-ROI connections survived multiple-comparison correction (Supplementary Fig. 2 and Supplementary Table 4). None of these connections involved the hippocampus.

### Hippocampal–MCC connectivity is associated with updating-related representational integration

To examine whether hippocampal–cortical connectivity was associated with updating-related representational reorganization, we computed two ΔW−B measures from ΔRDMupdate. One was based on within- and between-location object pairs defined according to the original map structure, whereas the other was based on pairs defined according to the updated map structure, paralleling the original- and updated-model RDM analyses described above. Each of these two ΔW−B measures was entered as the dependent variable in a separate linear regression model, with hippocampal–MCC functional connectivity as the predictor of interest and mean framewise displacement (mean FD) as a covariate (Fig. 4b; see Methods). Both analyses included 38 participants with valid hippocampal representational and connectivity data, and variance inflation factors (VIFs) were low in both models (value = 1.06), indicating no evidence of multicollinearity.

Hippocampal–MCC connectivity was significantly associated with integration of the updated map structure after controlling for mean FD (*b* = −0.008, 95% CI [−0.014, −0.002], *t*(35) = −2.72, *p* = 0.010, FDR-corrected *p* = 0.020; standardized *β* = −0.414; Fig. 4b and Supplementary Table 7). Specifically, stronger hippocampal–MCC connectivity was associated with more negative ΔW−B values, indicating greater representational integration of the updated map structure. In contrast, connectivity was not reliably associated with differentiation of the original map structure after controlling for mean FD (*b* = 0.003, 95% CI [−0.006, 0.011], *t*(35) = 0.67, *p* = 0.508, FDR-corrected *p* = 0.508; standardized *β* = 0.113; Fig. 4b and Supplementary Table 7).

To directly test whether hippocampal–MCC connectivity was differentially associated with the updated- and original-map ΔW−B measures, we computed partial correlations between connectivity and each measure while controlling for mean FD. These partial correlations closely paralleled the standardized regression coefficients reported above (β = −0.414 and 0.113, respectively). A bootstrap comparison with 5,000 iterations revealed a significant difference between the two partial correlations (*r*_updated = −0.418, *r*_original = 0.112, Δ*r* = −0.530, *p* = 0.013, 95% CI [−0.912, −0.100]).

Robustness of the hippocampal–MCC connectivity coefficient was further evaluated using three complementary analyses (see Methods). First, a two-tailed permutation test with 5,000 iterations provided additional support for the negative association between hippocampal–MCC connectivity and the updated-map ΔW−B measure, based on the empirical null distribution of the unstandardized connectivity coefficient (*p* = 0.010). Second, a 95% bias-corrected and accelerated (BCa) bootstrap confidence interval for the standardized connectivity coefficient, derived from 5,000 participant-level resamples, excluded zero (95% CI [−0.666, −0.093]). Third, in a leave-one-participant-out analysis, the unstandardized connectivity coefficient remained negative across all 38 iterations (range, −0.0092 to −0.0068; SD = 0.0005), indicating that the association was not disproportionately driven by any single participant.

Together, these results suggest that hippocampal–MCC connectivity was selectively associated with representational integration of the updated map, with no comparable evidence for an association with differentiation of the original map.

## Discussion

The present findings support a view of cognitive map updating as representational reorganization rather than complete overwriting of prior structure. In the right hippocampus, updated associations became relatively integrated while the prior relational structure remained differentiated, indicating that new and prior relational structures could coexist within hippocampal representational geometry. Map updating was accompanied by activation across frontoparietal, cingulate, and cerebellar regions, and right hippocampal–MCC connectivity was selectively associated with integration of the updated, but not the prior relational structure. Together, these results suggest that flexible cognitive map updating is accompanied by coordinated hippocampal–cortical interactions that incorporate new information while retaining prior relational structure.

### Hippocampal representational reorganization enables coexistence of prior and updated relational structures

Our results therefore suggest that hippocampal updating involves adaptive reorganization rather than wholesale replacement: prior relational structure is preserved while new information is incorporated through concurrent differentiation and integration. Following initial learning, objects sharing a location became more dissimilar, reflecting representational differentiation among overlapping associations—a process consistent with pattern separation-like mechanisms^35–37^. Following map updating, newly reassigned object–location pairs became more similar, indicating integration within a common representational context, in line with accounts proposing that reactivation of prior memories during new learning promotes integrative hippocampal coding^17,38^. Critically, this integration of the updated structure emerged without erasing the differentiation established during initial learning—the two processes coexisted within the right hippocampus, consistent with prior evidence of such coexistence^19,39^, and reflecting a balance between preserving prior knowledge and accommodating new information.

Notably, both model-based associations were attenuated after controlling for the alternative model RDM, suggesting that original- and updated-map representations were not fully independent but shared some representational variance. These findings raise the possibility that hippocampal updating operates through restructuring within a shared representational space, enabling new information to be incorporated while preserving prior relational structure. Such a representational format may support adaptive revision while minimizing catastrophic interference with existing knowledge^22,24,40^.

The lateralization of these effects warrants further consideration. Evidence for right hippocampal predominance was more apparent during initial learning than during map updating. This learning-related right hippocampal effect is broadly consistent with previous reports linking the right hippocampus to spatial relational processing and cognitive map formation^41,42^. In contrast, although updating-related effects were numerically stronger in the right hippocampus, representational reorganization was not significantly lateralized. This pattern raises the possibility that map updating, unlike initial map formation, imposes additional computational demands that may not be supported by strongly right-lateralized hippocampal mechanisms alone—a possibility consistent with the broader, more distributed cortical recruitment observed during updating.

### Distinct neural recruitment during cognitive map updating and maintenance

Beyond hippocampal representational reorganization, whole-brain analyses revealed that cognitive map updating and maintenance recruited largely distinct cortical systems, suggesting differences in their underlying computational demands.

Map updating preferentially recruited regions associated with attentional control and spatial processing, particularly the FEF, IPS, and SPL within the dorsal attention network, consistent with their established roles in spatial priority mapping^43^ and voluntary attentional orienting^44^. Activation also extended to the precuneus, a region implicated in visuospatial imagery and egocentric spatial updating^45,46^. Cerebellar activation was also observed during map updating (Fig. 3a). Given that motor demands were matched between shift and non-shift trials, this pattern more likely reflects cerebellar contributions to non-motor cognitive operations, consistent with established evidence for cerebellar involvement in higher-order cognition^47^.

In contrast, maintenance-related activation was greater in posterior occipital, lateral temporal, and angular gyrus regions associated with visual object processing, memory retrieval, and multimodal integration^48,49^. In the absence of relational restructuring demands, map maintenance may rely primarily on the retrieval and reinstatement of previously established object–location associations, rather than on the attentional and control processes engaged during updating.

Notably, both conditions recruited the midcingulate cortex, though activation was localized to anatomically and functionally distinct subregions. Updating-related activation was located in the anterior MCC bordering the pre-supplementary motor area, a region frequently implicated in conflict monitoring and cognitive control^50,51^, whereas maintenance-related activation was centered in a more posterior MCC region associated with response selection and motor control^52^. This spatial dissociation raises the possibility that map updating, relative to maintenance, recruits additional cognitive control resources to help resolve competition between newly updated and previously established representations.

### Hippocampal–MCC connectivity is selectively associated with integration of the updated map

Although representational analyses revealed hippocampal signatures of map updating, we did not observe significant hippocampal effects in the univariate analyses. Methodological work indicates that multivariate approaches are particularly sensitive to representational changes that may not be detectable using univariate mean signal differences^53^. Consistent with this view, the present findings suggest that hippocampal contributions to cognitive map updating may be more readily expressed through changes in representational geometry than through differences in overall activation magnitude. We next asked whether task-modulated hippocampal–cortical connectivity was associated with the degree of representational reorganization within the right hippocampus.

At the network level, cognitive map updating was characterized by widespread task-modulated functional connectivity, particularly among frontal and parietal regions. This pattern is consistent with distributed frontoparietal interactions supporting the flexible reorganization of previously learned relational knowledge^54,55^. Within this connectivity profile, the only significant hippocampal connection was observed between the right hippocampus and right MCC.

Hippocampal–MCC connectivity was selectively associated with integration of the updated relational structure, with no reliable association with differentiation of the prior relational structure. This selectivity suggests that hippocampal–MCC coupling may be more closely linked to incorporating updated associations than to general relational retrieval or maintenance. Because updated and prior associations were mapped onto the same set of spatial locations, integration may have required resolving competition between newly acquired and previously learned associations. This interpretation is broadly consistent with proposed roles of the MCC in monitoring and resolving competing task demands^50,51^.

Schema-based learning paradigms have implicated the ventromedial prefrontal cortex (vmPFC) in the rapid integration of schema-congruent information^56–58^, with vmPFC activity increasing as new information becomes more congruent with prior knowledge^59^. In contrast, the present findings point to MCC rather than vmPFC involvement, raising the possibility that distinct medial frontal regions contribute to memory updating under different computational demands. Specifically, vmPFC may be preferentially engaged when new information can be assimilated into existing knowledge structures, whereas MCC recruitment may become more prominent when updating requires resolving competition between incompatible relational representations. In the present task, updated object–location associations were established within the same spatial scaffold as the prior associations, creating competition between updated and previously learned relational structures. The present findings are broadly consistent with MCC involvement in competition-driven updating, insofar as hippocampal–MCC connectivity was selectively associated with integration of the updated relational structure, but not differentiation of the prior relational structure. Building on evidence linking hippocampal–posterior mPFC coupling to the integration of overlapping associative memories^28^, these findings raise the possibility that distinct medial frontal systems are recruited depending on whether updating is driven primarily by schema congruency or representational competition. However, the present design does not directly compare schema-congruent and competition-driven forms of updating. Future studies manipulating both factors within the same paradigm could clarify the conditions under which vmPFC and MCC are differentially engaged.

Nevertheless, these findings do not resolve the directionality of hippocampal–MCC communication. One possibility is that relational mismatch signals emerge within hippocampal representations and are transmitted to the MCC, engaging control processes that facilitate selective integration^25^. Alternatively, conflict-related signals originating in the MCC may modulate hippocampal updating in a demand-sensitive manner. Further research could address this question by examining the direction of information flow between the hippocampus and MCC using methods that combine high spatial resolution with high temporal sensitivity, such as intracranial recordings.

### Limitations and future directions

Although the present study provides insights into cognitive map updating, several limitations should be considered. First, the paradigm employed a minimal spatial scaffold comprising three discrete locations and nine objects, enabling precise experimental control over relational structure and its updating. However, this design differs from the richer environments typically used in classical cognitive map research, and it remains unclear whether the mechanisms identified here generalize to larger-scale settings involving continuous space and active navigation. Extending the present approach to more naturalistic spatial paradigms will therefore be an important next step.

Second, the findings were obtained over a relatively short three-day interval and may not capture the longer-term dynamics of map updating. Although both original and updated relational structures were detectable approximately one day after updating, the subsequent evolution of this representational configuration remains unknown. Systems consolidation^22^ may further alter the relative strength or organization of these representations over weeks or months. Future longitudinal studies tracking hippocampal representational geometry across longer delays could determine whether the original and updated structures continue to coexist or whether the updated structure gradually becomes predominant.

Third, the analyses used whole-hippocampus ROIs derived from 2-mm isotropic fMRI data, limiting the ability to resolve contributions from individual hippocampal subfields. Pattern separation and pattern completion computations are thought to depend on partially distinct circuitry involving the dentate gyrus, CA3, and CA1, and may therefore contribute differently to the preservation of prior structure and the integration of updated associations. Future studies using high-resolution hippocampal imaging could help clarify how distinct hippocampal subfields contribute to the coexistence and reorganization of relational map representations.

### Summary

Our findings identify a multilevel neural signature of cognitive map updating. The right hippocampus showed representational reorganization in which updated associations became integrated while prior relational structure remained differentiated. At the network level, map updating engaged frontoparietal control systems and was accompanied by hippocampal–midcingulate connectivity that was selectively associated with integration of the updated map structure, with no comparable evidence for differentiation of the original map structure. Together, these results suggest that cognitive map updating involves coordinated hippocampal–cortical processes that support the adaptive restructuring of relational knowledge without completely overwriting prior structure.

## Methods

### Participants

Forty-five healthy university students (mean age = 20.51 years, SD = 3.06, range = 18–28; 17 females) were recruited through the university’s online platform for a three-day experimental protocol. All participants had normal or corrected-to-normal vision and no history of neurological or psychiatric disorders. Written informed consent was obtained from all participants in accordance with the experimental protocol approved by the Ethics Committee of Kochi University of Technology, Japan. Participants were screened for MRI contraindications prior to enrollment and received 10,000 JPY as compensation. Sample size was guided by prior fMRI studies examining hippocampal representations and memory updating, which typically included approximately thirty participants^19,34^.

Of the forty-five recruited participants, all completed the spatial map learning task (behavioral session) on Day 1. Forty-four participants completed the spatial map updating task (fMRI session) on Day 2, as one participant withdrew because of physical discomfort. Forty-three participants completed the final picture viewing task (fMRI session) on Day 3, as two participants were absent from the final experimental session.

For the spatial map updating task, five additional participants were excluded from fMRI analyses due to excessive head movement (mean FD > 0.25 mm) or falling asleep during scanning, resulting in a final analytic sample of 39 participants. No participants were excluded from the picture viewing task based on these criteria, resulting in a final analytic sample of 43 participants for this task. Participant inclusion and exclusion across tasks and analyses are summarized in Table 1.

### Stimuli

Ten novel multicolored 3D-rendered objects were created using Blender 4.1 (www.blender.org)^17,60^. These stimuli were designed to be physically plausible yet distinct from real-world items to ensure that participants had no pre-existing representations. To control for low-level visual properties, all objects were equated for luminance and contrast using the SHINE toolbox in MATLAB (R2024a)^61^.

To provide a naturalistic visual context, we created a background consisting of a perspective grid with depth cues in Adobe Illustrator (Adobe Inc.). The grid was presented on a gray background (RGB: 160, 160, 160), together with three white square frames (3.5° × 3.5°). The frames were positioned at the upper center, lower left, and lower right of the screen (Supplementary Fig. 1a) and served as fixed spatial reference locations throughout the experiment. One of the ten objects was consistently designated as the target stimulus in all picture viewing tasks. The remaining nine objects were used in the spatial map learning task and were randomly assigned to one of the three spatial locations (UP, LEFT, or RIGHT) across participants.

### Paradigm overview

We implemented a three-day experimental paradigm to dissociate the formation and updating of cognitive maps (Fig. 1a). Detailed procedures for each task are described below.

### Picture Viewing Task (fMRI session)

The PVT was used to probe changes in neural pattern representations across learning and updating. In each session, participants viewed ten novel objects: nine task-relevant objects and one target object included to maintain attention. Participants were instructed to press a button whenever the target object appeared (Fig. 1b).

The task followed a mini-block design to ensure balanced exposure. Each mini-block included all ten objects presented in a pseudorandom order, with the constraint that no object appeared consecutively across block transitions. Each object was presented 12 times per session (2 runs, 6 mini-blocks per run), totaling 120 trials. In each trial, the stimulus was presented for 3 TRs (repetition time [TR] = 743 ms), followed by an inter-trial interval (ITI) of 5 or 7 TRs during which a fixation cross was displayed. The two interval durations occurred with equal probability.

The presentation order of the objects was randomized among participants but remained identical across the three sessions (PVT pre-learn, post-learn, and post-update) to minimize order-related confounding. Prior to the first scan, participants were familiarized with the stimuli to mitigate novelty-related neural responses and practiced the target-detection task. The target appeared in 10% of the trials, and these trials were excluded from subsequent multivariate pattern analyses. Behavioral performance on target trials was monitored to ensure alertness.

### Spatial Map Learning Task (behavioral session)

Following the PVT pre-learn scan, participants completed a spatial map learning task outside the scanner to establish associations between nine objects and three designated spatial locations (Fig. 1c).

Each trial commenced with a stimulus presented at the center of the screen for 2.23 s, followed by a 0.5 s fixation cross. Subsequently, three white square frames were displayed on a grid background, prompting participants to select the associated location using a response box. Upon each selection, 2 s of visual feedback was provided: a correct choice turned the square frame green and displayed the object within it, while an incorrect choice turned the frame red. After an incorrect response, participants were prompted to select again and continued until the correct location was identified. Each trial concluded with a 3 s ITI. The learning task consisted of five runs, with each object appearing twice per run. Stimuli were presented in a pseudorandomized mini-blocks to ensure all nine objects were encountered before any repetition occurred. On average, the task lasted approximately 15 min.

For each participant, the nine objects were assigned to the three locations, with three objects assigned to each location. Object–location mappings remained fixed throughout the learning task but were randomized across participants to minimize potential biases related to the visual or semantic properties of individual objects.

### Spatial Map Updating Task (fMRI session)

On Day 2, following the PVT post-learn scan, participants performed the spatial map updating task inside the scanner (Fig. 1d). The task required participants to update previously learned object–location associations in response to shift cues, consistent with prior work on cue-triggered representational updating^33^.

Three cue types were used: non-shift (NS), indicated by two semicircular arcs and instructing participants to retain the object’s original location; clockwise shift (CS), indicated by arcs with clockwise arrows and instructing participants to move the object’s associated location one position clockwise; and counterclockwise shift (CCS), indicated by arcs with counterclockwise arrows and instructing participants to move the object’s associated location one position counterclockwise. Cue assignment was balanced across the Day 1 spatial map: within each original location, the three associated objects were assigned one each to the CS, CCS, and NS conditions. Thus, each object was consistently associated with a single cue type, with three objects assigned to each condition, and no original location was systematically linked to a particular updating condition. Each object was presented 12 times across six runs, with two presentations per run, in pseudorandomized mini-blocks.

Each trial comprised four phases. During the retrieval phase, the object was presented at the center of the screen for 3 TRs, during which participants retrieved its original location. Following a 1-TR fixation interval, a shift cue was presented for 1 TR to indicate the transformation rule. During the subsequent delay phase, a fixation cross was displayed for 6 s, allowing participants to mentally update the object’s location while temporally separating rule processing from the motor response. In the response phase, three white square frames were presented on the grid background, and participants selected the updated location using an MRI-compatible response box. No feedback was provided, thereby minimizing reinforcement-related neural responses; instead, the selected frame remained visible for 0.5 s to acknowledge the response. Each trial concluded with a jittered ITI (5 or 7 TRs). The updating task lasted approximately 32 min in total.

### Memory assessment (behavioral session)

Object–location memory was assessed outside the scanner at four time points to track learning- and updating-related changes in spatial memory performance. Assessments were conducted immediately after the spatial map learning task on Day 1, following an overnight delay on Day 2, immediately after the spatial map updating task on Day 2, and following an overnight delay on Day 3. Importantly, the delayed assessments on Days 2 and 3 were administered only after the corresponding PVT session to minimize potential test-induced reactivation or cueing of the neural representations measured during the PVT.

On each trial, an object was presented, and participants indicated its associated location with a single response. No feedback was provided. Within each assessment, each of the nine objects was presented four times in a pseudorandom order, yielding 36 trials in total. Memory performance was quantified as the percentage of correct responses. Each assessment lasted approximately 3.5 min.

### fMRI data acquisition and preprocessing

Whole-brain imaging data were acquired on a Siemens Magnetom Prisma 3T MR scanner using a 64-channel head coil. A high-resolution T1-weighted anatomical image was acquired using an MPRAGE sequence (repetition time [TR] = 1900 ms; echo time [TE] = 2.52 ms; flip angle [FA] = 9°; field of view [FOV] = 250 mm; matrix = 256 × 256; in-plane resolution = 1 × 1 mm²; slice thickness = 1 mm; 176 slices). Functional images were collected using a multiband echo planar imaging (EPI) pulse sequence (TR = 743 ms; TE = 35.6 ms; FA = 48°; FOV = 192 mm; matrix = 96 × 96; in-plane resolution = 2 × 2 mm²; slice thickness = 2 mm; 72 slices; multiband acceleration factor = 8).

Anatomical and functional data were preprocessed with fMRIPrep (version 24.1.1, https://fmriprep.org/en/stable/)^62^, which is based on Nipype 1.8.6^63^.

### Anatomical data preprocessing

The T1-weighted (T1w) structural image was corrected for intensity non-uniformity (INU) using N4BiasFieldCorrection^64^ in ANTs 2.5.3^65^ and served as the T1w reference for the entire workflow. ANTs was then used to skull-strip the T1w-reference, resulting in the brain-extracted T1w. Brain tissue segmentation into cerebrospinal fluid (CSF), white matter (WM), and gray matter (GM) was performed using the FAST algorithm in FSL 6.0.7^66^. Brain surface reconstruction was conducted using recon-all in FreeSurfer 7.3.2^67^. Finally, the T1w image was spatially normalized to the MNI152NLin6Asym standard space (ICBM 152 Nonlinear Asymmetric template, version 2006)^68^. All anatomical outputs, including bias-corrected images, brain masks, and segmentations, were visually inspected via the HTML reports generated by fMRIPrep to ensure high-quality preprocessing.

### Functional data preprocessing

A reference BOLD volume was generated for head-motion estimation and correction using MCFLIRT (FSL 6.0.7). Functional images were co-registered to the T1-weighted anatomical reference using boundary-based registration (bbregister). The workflow resampled the BOLD time series into MNI152NLin6Asym standard space at 2 mm isotropic resolution.

Several nuisance regressors were estimated for quality control and subsequent modeling, including (1) motion parameters, their temporal derivatives, and quadratic terms; and (2) anatomical CompCor (aCompCor) components derived from cerebrospinal fluid (CSF) and white matter (WM) masks. These preprocessing steps followed the default fMRIPrep pipeline unless otherwise specified.

For different analytical purposes, the preprocessed data followed two distinct paths. For subsequent univariate activation analyses, motion-related noise components were first removed from the normalized BOLD time series using aggressive denoising based on Independent Component Analysis–based Automatic Removal of Motion Artifacts (ICA-AROMA)^69^. In contrast, representational similarity analyses were performed on unsmoothed BOLD time series in native space without ICA-AROMA denoising, to preserve fine-grained multivoxel activity patterns.

### Univariate general linear model analyses

Univariate GLM analysis was conducted on data acquired during the spatial map updating task using FSL (version 6.0.7.15). After exclusion (Table 1), a total of 39 participants were included in the univariate analysis. For each participant, first-level models included three task regressors corresponding to the CS, CCS, and NS conditions. Each cue was modeled as an event, time-locked to cue onset and lasting for 1 TR. Regressors were convolved with a canonical double-gamma hemodynamic response function (HRF), and temporal derivatives were included. Additional task events, including object presentation and manual responses, were modeled as nuisance regressors. Prior to model estimation, a high-pass temporal filter (100 s) was applied to remove low-frequency drifts, and the denoised fMRI data were smoothed using a Gaussian kernel (FWHM = 6 mm) to increase sensitivity. Time series autocorrelation was accounted for using FILM with local autocorrelation correction, as implemented in FSL.

For group-level inference, one-sample *t*-tests with 10,000 permutations were performed on the contrast images of shift trials (CS and CCS combined) relative to non-shift trials (NS) across all 39 participants. These analyses were implemented using the LISA framework^70^ within the LIPSIA package (version 3.1.0; https://github.com/lipsia-fmri/lipsia). LISA is a non-parametric and threshold-free approach that integrates spatial information via bilateral filtering, providing a sensitive and robust method for group-level analysis^71^. Multiple-comparison correction was applied using False Discovery Rate (FDR) at *p* < 0.01. A stringent FDR threshold was applied to reduce the risk of false positives and enhance the specificity of the reported results. All statistical maps were restricted to a group-level gray matter mask, defined as voxels with a mean gray matter probability greater than 10% across the entire sample.

### Representational similarity analysis (RSA)

Representational similarity analysis was conducted on fMRI data acquired during the PVT sessions to quantify neural representational changes associated with spatial map learning and updating. Analyses were performed separately for the PVT pre-learn, PVT post-learn, and PVT post-update sessions. As described in Table 1, two participants were absent from the final session, resulting in a final RSA sample of 43 participants.

### Item-level beta estimation

Prior to statistical modeling, the first six TRs of each functional run were discarded to allow for signal stabilization. For each run within each PVT session, voxel-wise GLMs were constructed using FSL (version 6.0.7). Each object was modeled with a separate regressor time-locked to stimulus onset to obtain item-level beta estimates. The design matrix included 10 regressors of interest: nine corresponding to task-relevant objects and one corresponding to the target object. Events were modeled as epochs lasting the duration of stimulus presentation (3 TRs) and convolved with a canonical HRF. Target trials were included in the GLM but excluded from subsequent RSA.

Nuisance regressors derived from fMRIPrep included six rigid-body motion parameters, framewise displacement, and six anatomical CompCor components extracted from CSF and WM masks. Separate nuisance regressors were additionally included for volumes with framewise displacement greater than 0.3 mm. Temporal high-pass filtering with a 100-s cutoff was applied within the FSL FEAT framework. To preserve fine-grained multivoxel information, GLMs were estimated using unsmoothed functional data. Item-level beta patterns for the nine task-relevant objects were subsequently entered into ROI-based RSA.

### ROI definition

Based on prior work on cognitive map formation in humans^72,8^, analyses were conducted separately for the left and right hippocampus. The hippocampal ROIs were defined using automated anatomical segmentation in FreeSurfer (version 7.3.2) on each participant’s high-resolution T1-weighted anatomical image, with masks extracted from the *aparc.DKTatlas+aseg* segmentation. Label maps were converted to binary masks and resampled into each participant’s native functional space using nearest-neighbor interpolation, with the first-level contrast image serving as the reference to ensure spatial alignment with the beta estimates used in subsequent analyses. All segmentations were visually inspected to ensure accurate anatomical localization.

### Crossnobis-based within–between dissimilarity analysis

Learning- and updating-related representational changes were quantified using cross-validated Mahalanobis distances. For each participant, ROI, phase, and run, condition-specific multivoxel patterns were extracted from the item-level GLM beta estimates. Noise covariance was estimated from residual time series, regularized using a shrinkage parameter (λ = 0.1) following prior crossnobis RSA implementations^73^, and used to whiten multivoxel activation patterns.

Cross-validated Mahalanobis distances were computed between all condition pairs using independent runs, yielding unbiased estimates of representational dissimilarity. Under cross-validation, distance estimates are centered around zero under the null hypothesis. The resulting values formed a representational dissimilarity matrix (RDM) for each participant, ROI, and phase.

Learning-related representational change was quantified as a neural ΔRDMlearn (RDM post-learn minus RDM pre-learn; Fig. 2a). Updating-related representational change was quantified as a neural ΔRDMupdate (RDM post-update minus RDM pre-learn; Fig. 2a). Pre-learning was used as the baseline for both comparisons, such that updating-related ΔRDMs reflected the cumulative representational changes associated with learning and subsequent map updating.

To quantify representational reorganization, pairwise dissimilarities were grouped as within-location or between-location object pairs. Within-location pairs were defined as object pairs occupying the same location under the relevant map structure, whereas between-location pairs comprised object pairs occupying different locations. Mean within-location and between-location dissimilarity changes were computed from the corresponding ΔRDMs, and a differentiation index (ΔW−B) was defined as the mean within-location change minus the mean between-location change. Positive ΔW−B values indicated greater increases in dissimilarity for within-location than between-location object pairs relative to baseline, consistent with representational differentiation. Conversely, negative ΔW−B values indicated relatively greater similarity among within-location object pairs, consistent with representational integration.

Statistical significance of ΔW−B values was assessed using one-sample *t*-tests across participants (*n* = 43). Values of *p* were corrected for multiple comparisons using Benjamini–Hochberg FDR correction (*p* < 0.05), applied separately within learning-related and updating-related analyses, with each family comprising tests across all ROIs (left and right hippocampus).

### Model-based representational similarity analysis

Complementing the within–between dissimilarity analysis described above, model-based RSA was performed to test whether learning- and updating-related neural representational changes conformed to the relational structure of the original and updated spatial maps. The neural RDMs used in this analysis were the same phase-specific crossnobis distance matrices generated for the within–between dissimilarity analysis. For each participant and hippocampal ROI, learning-related representational change was quantified as a neural ΔRDMlearn, and updating-related representational change was quantified as a neural ΔRDMupdate.

The original model RDM assigned a value of 1 to object pairs occupying the same location in the original map and 0 to pairs occupying different locations. The updated model RDM used the corresponding coding based on the updated map. These model RDMs therefore functioned as relational contrast models rather than conventional dissimilarity models. Diagonal elements were excluded, and the upper-triangular elements of the neural and model RDMs were vectorized. Spearman rank correlations were computed between the vectorized neural ΔRDM and the corresponding vectorized model RDMs. For the learning-related analysis, only the original model RDM was evaluated because the updated map had not yet been introduced. For the updating-related analysis, both the original and updated model RDMs were evaluated.

Because the model RDMs coded within-location pairs as 1 and between-location pairs as 0, positive correlations indicated a relative increase in neural dissimilarity among within-location compared with between-location pairs, consistent with representational differentiation. Conversely, negative correlations indicated a relative decrease in neural dissimilarity among within-location compared with between-location pairs, consistent with representational integration.

As a complementary analysis addressing the partial dependence between the original and updated model RDMs (Spearman’s ρ = −0.33), corresponding to approximately 11% shared variance), partial Spearman correlations were calculated for ΔRDMupdate. The original-model fit was estimated while controlling for the updated model RDM, and the updated-model fit was estimated while controlling for the original model RDM.

Correlation coefficients were Fisher *z*-transformed prior to group-level statistical testing. Statistical significance was assessed using one-sample *t*-tests across participants on Fisher *z*-transformed model fits. Benjamini–Hochberg FDR correction was applied across the left- and right-hippocampal tests separately for each phase-by-model comparison. All analyses were implemented in MATLAB (R2024a) using custom scripts.

### Task-based functional connectivity with gPPI

Generalized psychophysiological interaction^74^ implemented in the CONN toolbox was used to explore the functional connectivity between map updating-related regions.

### Preprocessing for gPPI analyses

Preprocessed fMRI data (fMRIPrep) were imported into the CONN toolbox for further analysis. To eliminate potential confounding effects, linear regression was applied, followed by high-pass filtering with a cutoff frequency of 0.008 Hz. The CONN toolbox’s predefined confound removal strategy incorporated several sources of noise, such as anatomical artifacts, motion-related distortions, and constant task-related signals^75^. Anatomical noise was composed of five principal components of WM and five principal components of CSF, which were calculated using aCompCor^76^. Motion artifacts were taken into account by including six head-motion parameters, first-order temporal derivatives, and time frames with excessive motion^77^. These anatomical and motion-related confounds were estimated with fMRIPrep.

Furthermore, task-related effects were modeled using standard task regressors convolved with a canonical double-gamma hemodynamic response function, along with their first-order temporal derivatives.

### ROI-to-ROI gPPI analyses in CONN

In the ROI-to-ROI gPPI analyses, excessive smoothing can inadvertently include signals from unrelated voxels, which can distort the properties of functional networks, including the node degrees^78^. To mitigate this potential issue, spatial smoothing was intentionally omitted here. In the current analysis, we employed a multiple regression approach to analyze functional connectivity across ROIs. The average BOLD signal of each ROI was used as a physiological regressor. For each pair of ROIs, a separate regression model was constructed, where the BOLD signal of the target ROI *R_j_*(*t*) was regressed on the BOLD signal of the seed ROI *R_i_*(*t*), along with psychological regressors representing the task conditions. The model is described by the equation:

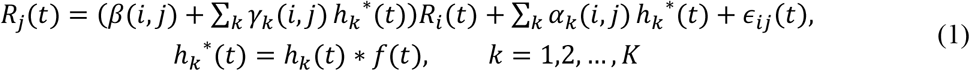

where the *k*-th psychological regressor, 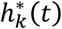 is obtained by convolving the task onset information ℎ*_k_*(*t*) with a canonical hemodynamic response function *f*(*t*). The optimal values of *α*, *β*, and *γ* were estimated by minimizing the residuals *ε_ij_*(*t*) using an ordinary least squares (OLS) solution:

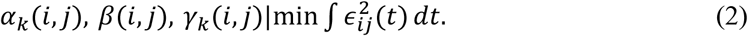

The parameter γ_k_(i, j) represents directed functional connectivity from the seed region *R_i_* to the target region R_j_ under the *k*-th psychological condition 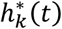. In contrast, *γ_k_*(*j*, *i*) represents directed functional connectivity from *R_j_* to R_i_. Our focus was on the functional connectivity between regions, rather than directional effective connectivity. Therefore, the functional connectivity between regions *R_i_* and *R_j_* was defined as the average of *γ_k_*(*i*, *j*) and *γ_k_*(*j*, *i*).

### gPPI analyses of map-updating and map-maintenance networks

To define task-related nodes for connectivity analyses, local activation peaks were extracted from the group-level statistical maps described above (LISA framework, FDR-corrected p < 0.01). Thirty-two peak coordinates from the shift > non-shift contrast were used to define the map-updating network, whereas twenty-four peak coordinates from the non-shift > shift contrast were used to define the map-maintenance network. Spherical ROIs with a 6-mm radius were centered on each peak. An 18-mm minimum spacing criterion—exceeding the sum of the sphere radii (12 mm)—was adopted to reduce spatial dependence and minimize overlap between neighboring functional nodes. Detailed MNI coordinates and associated statistics are provided in Supplementary Table 2. In addition to these activation-defined cortical ROIs, anatomically defined left and right hippocampal ROIs derived from FreeSurfer segmentations were included as a priori regions of interest.

At the group level, subject-specific gPPI interaction parameters were extracted from the CONN toolbox and imported into R (version 4.1.1) for ROI-to-ROI analyses. Functional connectivity between each ROI pair was defined as the average of *γ_k_*(*i*, *j*) and *γ_k_*(*j*, *i*), yielding a symmetric connectivity estimate. Statistical significance of task-modulated connectivity was assessed using two-tailed one-sample *t*-tests across participants. Multiple comparisons were controlled using the fdrtool package (version 1.2.18), which estimates the proportion of true null hypotheses and computes tail-area based FDR values (*p* < 0.05)^79,80^.

Because cortical ROIs were defined from task-related activation contrasts in the same sample, connectivity results involving these regions were interpreted as characterizing interactions within the task-responsive networks, rather than as fully independent evidence for the involvement of specific cortical regions. In contrast, hippocampal ROIs were anatomically defined a priori and were therefore independent of the task activation contrasts used for cortical ROI selection.

### Connectivity–representation association analyses

To examine whether hippocampal–cortical functional connectivity during updating was associated with representational reorganization, we related representational reorganization measures to task-modulated connectivity estimates obtained from significant hippocampal–midcingulate connectivity identified in the gPPI analysis. Two representational reorganization measures (ΔW−B) were computed from the same updating-related ΔRDM (post-updating minus pre-learning), by classifying object pairs as within- or between-location according to either the original map structure or the updated map structure, yielding ΔW−B_original and ΔW−B_updated, respectively. Analyses were restricted to 38 participants with valid PVT-derived representational measures, updating-task gPPI estimates, and motion estimates.

Two ordinary least-squares multiple linear regression models were estimated in R (version 4.1.1), one for each representational measure:

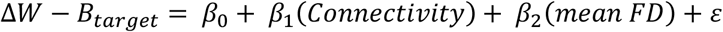

where Δ*W* − *B_target_* refers to either ΔW−B_original or ΔW−B_updated. Hippocampal–MCC connectivity was entered as the predictor of interest, and mean FD was included as a nuisance covariate. The *p*-values for the hippocampal–MCC connectivity coefficients were corrected across the updated- and original-map regression models using the Benjamini–Hochberg false discovery rate procedure. Variance inflation factors were examined, with values below 5 taken to indicate no substantial multicollinearity. Standardized regression coefficients were obtained by *z*-scoring all variables prior to model estimation, with variables re-standardized within each bootstrap sample during robustness checks.

To directly test whether hippocampal–MCC connectivity was differentially associated with the updated and original map structures, Pearson partial correlations were computed between connectivity and each ΔW−B measure while controlling for mean FD. Partial correlations were obtained by separately residualizing connectivity and each representational measure with respect to mean FD and correlating the resulting residuals. The difference between the two partial correlations, Δ*r* = *r*_updated − *r*_original, was evaluated using 5,000 paired participant-level bootstrap resamples. Within each resample, participants were sampled with replacement using the same resampling indices for all variables, and the residualization and correlation procedures were repeated. A percentile-based 95% confidence interval was obtained from the 2.5th and 97.5th percentiles of the bootstrap distribution of Δ*r*. For the two-tailed bootstrap *p* value, the bootstrap distribution was centered by subtracting its mean, and the *p* value was calculated as the proportion of centered bootstrap differences whose absolute values were greater than or equal to the absolute observed Δ*r*.

Robustness of the hippocampal–MCC connectivity coefficient in the updated-map model was assessed using three complementary analyses. First, a permutation test with 5,000 iterations was performed by randomly permuting connectivity values across participants while retaining the outcome and mean FD in their original order. The regression model was refitted after each permutation, and the observed unstandardized connectivity coefficient was compared with the resulting empirical null distribution using a two-tailed test. Second, a 95% bias-corrected and accelerated bootstrap confidence interval for the standardized connectivity coefficient was obtained from 5,000 participant-level resamples. The outcome, connectivity, and mean FD were re-standardized within each bootstrap sample before model estimation.

Third, a descriptive leave-one-participant-out analysis was conducted by iteratively refitting the model after excluding one participant at a time, yielding one coefficient estimate per omitted participant. The sign and range of these estimates were then examined.

## Code availability

Analysis code is available on the Open Science Framework.

## Author contributions

J.F. and K.N. conceived the project. J.F., C.H., and K.N. collected data. J.F., C.H. and R.W. analyzed data. J.F., R.W., I.H., K.J. and K.N. wrote the manuscript.

## Acknowledgments

We would like to thank Ms. Maoko Yamanaka for her administrative assistance. This study was supported by KAKENHI from Japan Society for the Promotion of Science (23H00413 to K.N. and I.H), and by AMED under grant number JP26wm0625205 to K.N. and I.H.

## Supplemental Material

**Supplementary Fig. 1:**
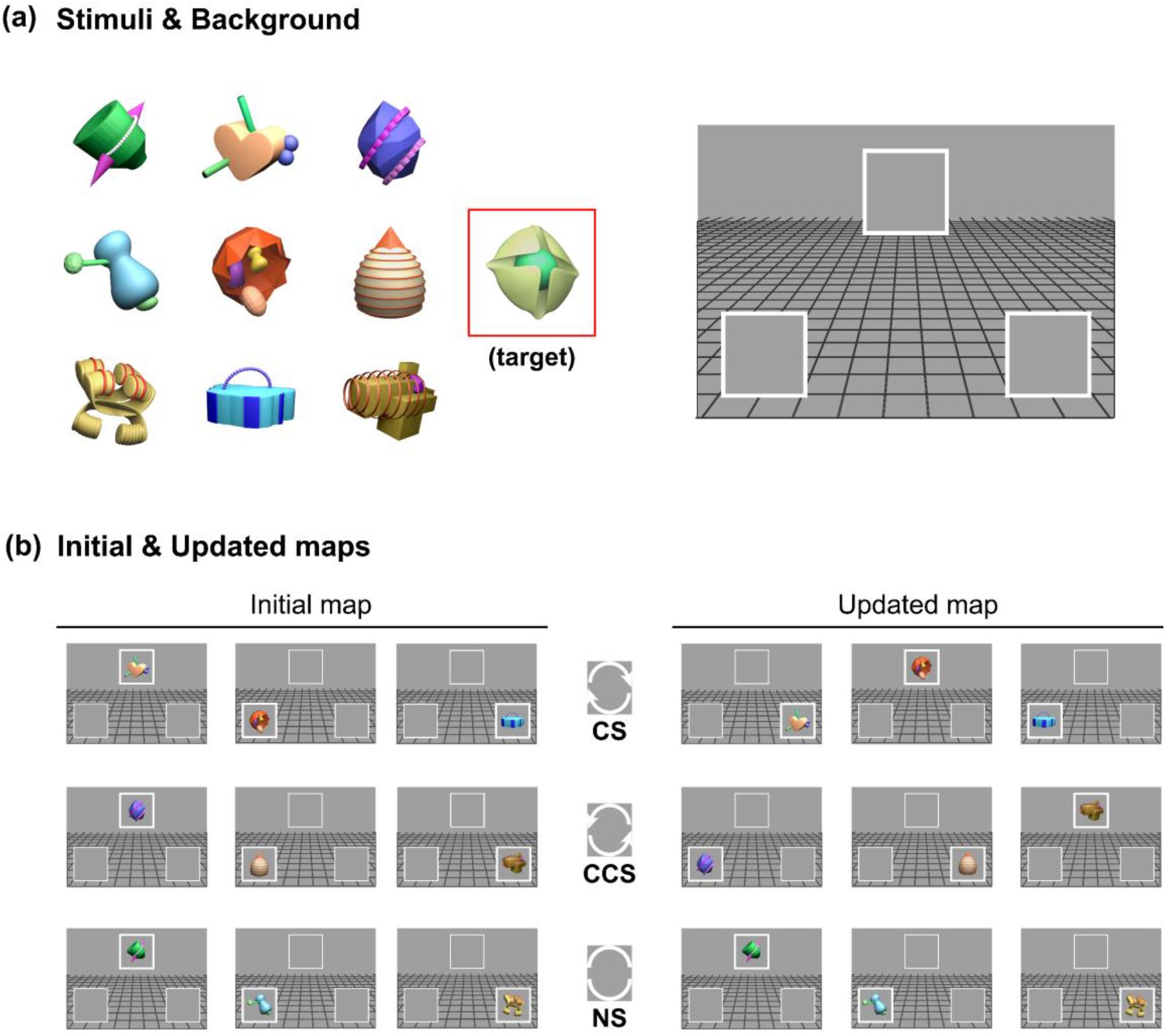
Illustration of novel stimuli and the schematic for cognitive map formation and updating. (a) Stimuli and background. The stimulus set consisted of ten novel multicolored 3D-rendered objects. All ten objects were presented in the picture viewing task, with one object serving as the target. The remaining nine objects were used in the spatial map learning and updating tasks. The background display consisted of a grid extending from the bottom of the screen toward the horizon, with three possible goal locations indicated by white square frames. (b) Initial and updated spatial maps. Left: initial map established after learning, in which participants learned associations between nine novel objects and three designated goal locations. Right: updated map following the updating task, in which participants revised the previously learned object–location associations according to cue-defined spatial transformations. The middle arrows illustrate the three possible transformations: clockwise shift (CS), counterclockwise shift (CCS), and non-shift (NS). Each object was consistently paired with one of these cue types during the updating task.

**Supplementary Fig. 2:**
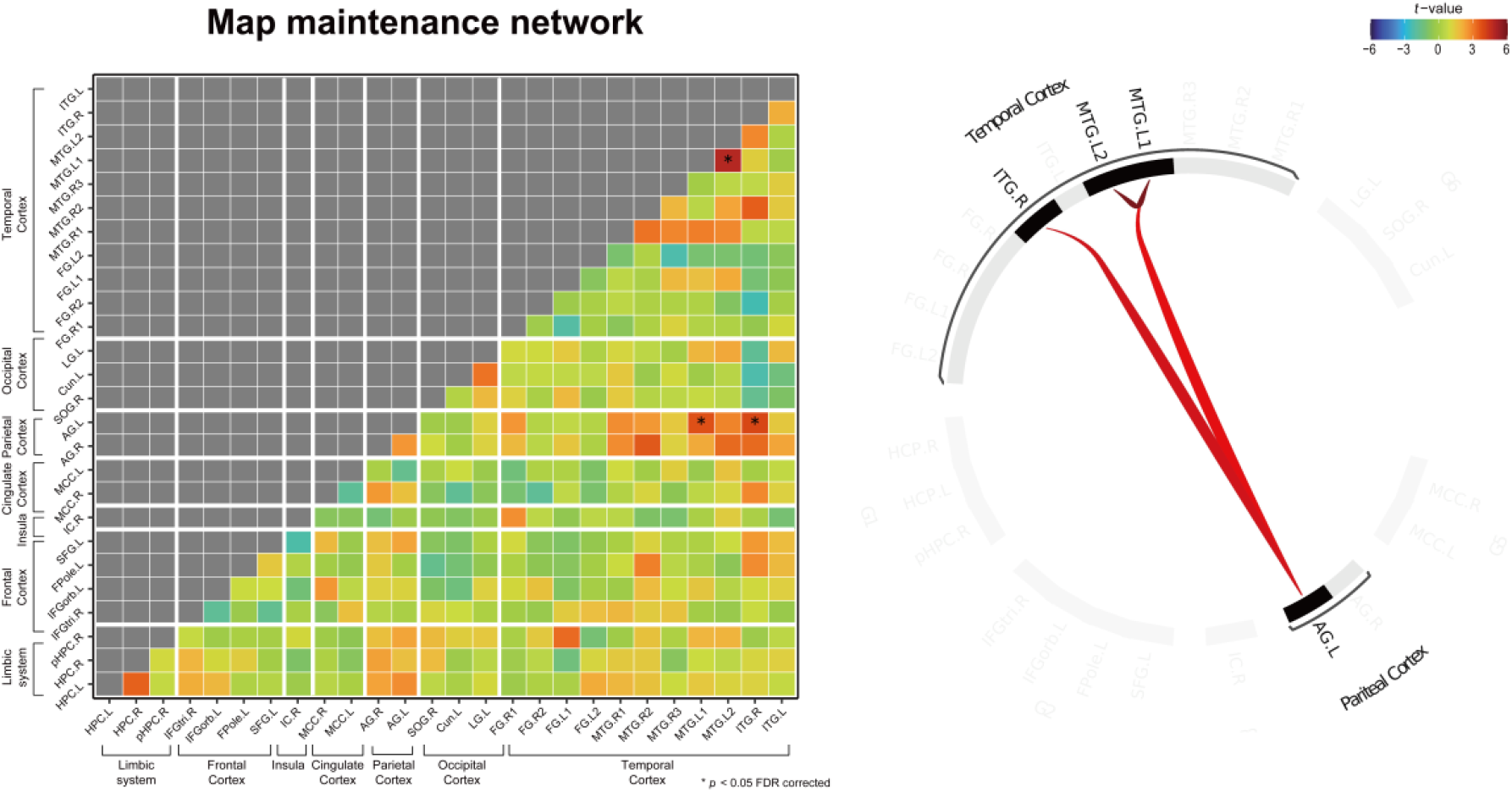
Task-modulated functional connectivity within the map maintenance-related network. The heatmap (left) illustrates task-modulated functional connectivity among regions identified by the map maintenance-related contrast (non-shift > shift), with color intensity reflecting *t*-values. The connectogram (right) is thresholded to highlight significant functional connections (*p* < 0.05, FDR-corrected; marked by asterisks in the heatmap). The maintenance-related network exhibited a sparse connectivity profile, with significant interactions confined to temporal and parietal regions. Specifically, the left angular gyrus (AG.L) showed significant connectivity with two left middle temporal gyrus clusters (MTG.L1/MTG.L2) and the right inferior temporal gyrus (ITG.R). No significant task-modulated functional connectivity was identified for the hippocampus in either hemisphere.

**Supplementary Table 1.** Learning- and Updating-driven Representational Changes in Hippocampal ROIs.

| Hippocampal ROI | $\Delta W-B$ | $t(42)$ | $p$ | $p_{\text{FDR}}$<br>(Benjamini-Hochberg) | Cohen's $d$ |
| --- | --- | --- | --- | --- | --- |
| **PVT post-learn minus PVT pre-learn** |  |  |  |  |  |
| HPC.L | 0.0013 | 0.159 | 0.874 | 0.874 | 0.024 |
| HPC.R | 0.0059 | 2.800 | 0.008 | 0.015 | 0.427 |
| **PVT post-update minus PVT pre-learn (original map)** |  |  |  |  |  |
| HPC.L | -0.0011 | -0.464 | 0.645 | 0.645 | -0.071 |
| HPC.R | 0.0050 | 2.636 | 0.012 | 0.023 | 0.402 |
| **PVT post-update minus PVT pre-learn (updated map)** |  |  |  |  |  |
| HPC.L | -0.0022 | -1.133 | 0.264 | 0.264 | -0.173 |
| HPC.R | -0.0053 | -3.364 | 0.002 | 0.003 | -0.513 |

*Note.* The within-between dissimilarity change index ( $\Delta W-B$ ) was calculated for the left and right hippocampus (HPC; L = left, R = right) across the PVT session contrasts. $\Delta W-B$ represents the change in representational dissimilarity for within-location minus between-location object pairs. Positive values indicate representational differentiation, whereas negative values indicate representational integration.

**Supplementary Table 2.** Coordinates and statistical properties of task-defined ROIs.

| Cluster | Size<br>(voxels) | ROI<br>id | Region | Full Name (Hemisphere) | Peak<br>$p$_FDR | Peak MNI<br>Coordinates |
| --- | --- | --- | --- | --- | --- | --- |
| ***shift > non-shift*** |  |  |  |  |  |  |
| C1 | 12392 | 1 | SPL.L | Superior Parietal Lobule (L) | < 0.001 | -28, -56, 50 |
| C1 | 12392 | 2 | PCun.R1 | Precuneus (R) | < 0.001 | 14, -68, 50 |
| C1 | 12392 | 3 | PCun.L1 | Precuneus (L) | < 0.001 | -10, -74, 50 |
| C1 | 12392 | 4 | PreCG.L | Precentral Gyrus (L) | < 0.001 | -20, -10, 48 |
| C1 | 12392 | 5 | IPL.L1 | Inferior Parietal Lobule (L) | < 0.001 | -50, -26, 46 |
| C1 | 12392 | 6 | IPS.L1 | Intraparietal Sulcus (L) | < 0.001 | -34, -44, 34 |
| C2 | 1984 | 7 | CereVI.R1 | Cerebellum Lobule VI (R) | < 0.001 | 26, -54, -24 |
| C2 | 1984 | 8 | CereVIII.R | Cerebellum Lobule VIII (R) | < 0.001 | 26, -52, -46 |
| C2 | 1984 | 9 | CereVI.R2 | Cerebellum Lobule VI (R) | < 0.001 | 34, -38, -34 |
| C2 | 1984 | 10 | Vermis6 | Vermis lobule VI | < 0.001 | 6, -66, -12 |
| C2 | 1984 | 11 | Vermis7 | Vermis lobule VII | < 0.001 | 4, -78, -26 |
| C3 | 1375 | 12 | PreCG.R1 | Precentral Gyrus (R) | < 0.001 | 28, -8, 46 |
| C4 | 1079 | 13 | SMA.L | Supplementary Motor Area (L) | < 0.001 | -4, 12, 50 |
| C4 | 1079 | 14 | MCC.R | Midcingulate Cortex (R) | < 0.001 | 14, 16, 36 |
| C5 | 423 | 15 | CereVII.L | Cerebellum Lobule VII (L) | < 0.001 | -38, -38, -32 |
| C5 | 423 | 16 | CereVIII.L | Cerebellum Lobule VIII (L) | < 0.001 | -34, -40, -50 |
| C5 | 423 | 17 | CereX.L | Cerebellum Lobule X (L) | < 0.001 | -18, -34, -42 |
| C6 | 352 | 18 | IPS.L2 | Intraparietal Sulcus (L) | < 0.001 | -30, -66, 26 |
| C6 | 352 | 19 | IPL.L2 | Inferior Parietal Lobule (L) | < 0.001 | -28, -84, 42 |
| C7 | 338 | 20 | Put.L1 | Putamen (L) | < 0.001 | -28, -18, -2 |
| C7 | 338 | 21 | Put.L2 | Putamen (L) | < 0.001 | -22, 12, 8 |
| C8 | 204 | 22 | PCun.R2 | Precuneus (R) | < 0.001 | 18, -58, 22 |
| C9 | 199 | 23 | IC.L | Insular Cortex (L) | < 0.001 | -28, 28, 10 |
| C10 | 141 | 24 | IPS.R | Intraparietal Sulcus (R) | 0.001 | 40, -74, 32 |
| C11 | 134 | 25 | Cun.L | Cuneus (L) | 0.001 | -14, -66, 24 |
| C12 | 90 | 26 | Thal.L | Thalamus (L) | < 0.001 | -16, -24, 10 |
| C13 | 54 | 27 | PreCG.R2 | Precentral Gyrus (R) | 0.003 | 62, 10, 30 |
| C14 | 52 | 28 | CereVI.L | Cerebellum Lobule VI (L) | 0.002 | -24, -58, -24 |
| C15 | 44 | 29 | MOG.L | Middle Occipital Gyrus (L) | 0.001 | -48, -80, -2 |
| C16 | 41 | 30 | Put.R | Putamen (R) | 0.002 | 22, 18, 2 |
| C17 | 36 | 31 | IC.R | Insular Cortex (R) | 0.004 | 32, 26, 2 |
| C18 | 32 | 32 | Vermis9 | Vermis lobule X | 0.003 | -2, -58, -34 |
| ***non-shift > shift*** |  |  |  |  |  |  |
| C1 | 1000 | 1 | SOG.R | Superior Occipital Gyrus (R) | < 0.001 | 14, -96, 20 |
| C2 | 955 | 2 | MCC.R | Midcingulate Cortex (R) | < 0.001 | 4, -44, 34 |
| C2 | 955 | 3 | MCC.L | Midcingulate Cortex (L) | 0.002 | -6, -20, 42 |
| C3 | 741 | 4 | Cun.L | Cuneus (L) | < 0.001 | -8, -92, 20 |
| C4 | 606 | 5 | FG.R1 | Fusiform Gyrus (R) | < 0.001 | 30, -52, -6 |
| C4 | 606 | 6 | FG.R2 | Fusiform Gyrus (R) | 0.001 | 30, -70, -10 |
| C5 | 478 | 7 | ITG.R | Inferior Temporal Gyrus (R) | < 0.001 | 56, -8, -32 |
| C6 | 419 | 8 | MTG.R1 | Middle Temporal Gyrus (R) | 0.001 | 52, -16, -10 |
| C6 | 419 | 9 | MTG.R2 | Middle Temporal Gyrus (R) | 0.001 | 68, -44, 4 |
| C6 | 419 | 10 | MTG.R3 | Middle Temporal Gyrus (R) | 0.005 | 52, -36, -4 |
| C7 | 283 | 11 | AG.R | Angular Gyrus (R) | 0.001 | 46, -54, 24 |
| C8 | 266 | 12 | FG.L1 | Fusiform Gyrus (L) | < 0.001 | -32, -50, -18 |
| C8 | 266 | 13 | FG.L2 | Fusiform Gyrus (L) | 0.006 | -22, -34, -16 |
| C9 | 187 | 14 | IFGtri.R | Inferior Frontal Gyrus, pars triangularis (R) | 0.002 | 56, 38, 8 |
| C10 | 126 | 15 | MTG.L1 | Middle Temporal Gyrus (L) | 0.002 | -58, -34, -2 |
| C11 | 88 | 16 | LG.L | Lingual Gyrus (L) | 0.005 | -22, -66, -10 |
| C12 | 77 | 17 | AG.L | Angular Gyrus (L) | 0.002 | -48, -62, 28 |
| C13 | 71 | 18 | ParaHPC.R | Parahippocampal Gyrus (R) | 0.002 | 36, -12, -28 |
| C14 | 55 | 19 | IFGorb.L | Inferior Frontal Gyrus, pars orbitalis (L) | 0.003 | -32, 38, -8 |
| C15 | 46 | 20 | FPole.L | Frontal Pole (L) | 0.004 | 8, 60, 12 |
| C16 | 44 | 21 | ITG.L | Inferior Temporal Gyrus (L) | 0.004 | -56, -10, -30 |
| C17 | 42 | 22 | MTG.L2 | Middle Temporal Gyrus (L) | 0.005 | -58, -18, -6 |
| C18 | 41 | 23 | IC.R | Insular Cortex (R) | 0.002 | 44, -2, -4 |
| C19 | 33 | 24 | SFG.L | Superior Frontal Gyrus (L) | 0.003 | -16, 30, 62 |

*Note.* Seed regions were derived from whole-brain univariate activation maps for the shift > non-shift and non-shift > shift contrasts, thresholded using voxel-level FDR correction with a minimum cluster size of 30 voxels. Within each significant cluster, local maxima were identified and used to define spherical ROIs with a 6-mm radius for subsequent analyses. When multiple local maxima were present within the same anatomical region, peaks were retained only if they were separated by at least 18 mm; otherwise, the peak with the highest statistical significance was retained to avoid redundant comparisons. All coordinates are reported in MNI space.

**Supplementary Table 3.** Significant functional connectivity within the map updating network (FDR-corrected *p* < 0.05).

| No. | Regions | Connectivity Strength | $t(38)$ | $p$ | $p_{\text{FDR}}$ |
| --- | --- | --- | --- | --- | --- |
| 1 | **MCC.R–HPC.R** | **0.209** | **2.826** | **0.007** | **0.034** |
| 2 | PCun.L–IPS.L1 | 0.610 | 7.063 | < 0.001 | < 0.001 |
| 3 | PCun.R1–IPS.L1 | 0.568 | 6.940 | < 0.001 | < 0.001 |
| 4 | PCun.R1–IPS.R | 0.534 | 6.942 | < 0.001 | < 0.001 |
| 5 | PCun.R1–PCun.L | 0.718 | 6.410 | < 0.001 | < 0.001 |
| 6 | PCun.L–SPL.L | 0.675 | 6.409 | < 0.001 | < 0.001 |
| 7 | IPS.R–IPS.L1 | 0.452 | 5.834 | < 0.001 | < 0.001 |
| 8 | Vermis7–PCun.L | 0.430 | 5.723 | < 0.001 | < 0.001 |
| 9 | PCun.L–IPS.R | 0.711 | 5.605 | < 0.001 | < 0.001 |
| 10 | Cun.L–SMA.L | 0.452 | 5.581 | < 0.001 | < 0.001 |
| 11 | IPS.R–PreCG.R1 | 0.501 | 5.549 | < 0.001 | < 0.001 |
| 12 | IC.R–SMA.L | 0.467 | 5.494 | < 0.001 | < 0.001 |
| 13 | Cun.L–IC.R | 0.468 | 5.476 | < 0.001 | < 0.001 |
| 14 | IPS.L2–IPS.L1 | 0.359 | 5.473 | < 0.001 | < 0.001 |
| 15 | SPL.L–IPS.L1 | 0.468 | 5.245 | < 0.001 | < 0.001 |
| 16 | PCun.L–IPL.L2 | 0.627 | 5.162 | < 0.001 | < 0.001 |
| 17 | Cun.L–IPS.L1 | 0.424 | 5.015 | < 0.001 | < 0.001 |
| 18 | MOG.L–IPL.L2 | 0.490 | 4.978 | < 0.001 | < 0.001 |
| 19 | IPL.L1–PreCG.R1 | 0.381 | 4.833 | < 0.001 | < 0.001 |
| 20 | IPS.R–PreCG.L | 0.368 | 4.813 | < 0.001 | < 0.001 |
| 21 | CereX.L–SMA.L | 0.260 | 4.806 | < 0.001 | < 0.001 |
| 22 | PreCG.R2–PreCG.R1 | 0.409 | 4.674 | < 0.001 | 0.001 |
| 23 | PCun.R2–SMA.L | 0.410 | 4.667 | < 0.001 | 0.001 |
| 24 | SPL.L–IPS.R | 0.517 | 4.651 | < 0.001 | 0.001 |
| 25 | PCun.R2–PCun.R1 | 0.410 | 4.556 | < 0.001 | 0.001 |
| 26 | PCun.R2–IC.R | 0.379 | 4.531 | < 0.001 | 0.001 |
| 27 | PCun.R2–PCun.L | 0.445 | 4.424 | < 0.001 | 0.001 |
| 28 | PCun.R1–SPL.L | 0.514 | 4.408 | < 0.001 | 0.001 |
| 29 | PCun.R1–PreCG.R1 | 0.406 | 4.246 | < 0.001 | 0.002 |
| 30 | IPS.R–IC.R | 0.387 | 4.227 | < 0.001 | 0.002 |
| 31 | IPL.L1–PreCG.R2 | 0.432 | 4.206 | < 0.001 | 0.002 |
| 32 | PCun.R1–IC.L | 0.276 | 4.151 | < 0.001 | 0.002 |
| 33 | PCun.R2–IPS.L1 | 0.340 | 4.116 | < 0.001 | 0.002 |
| 34 | Vermis7–IPL.L2 | 0.342 | 4.099 | < 0.001 | 0.002 |
| 35 | PCun.L–PreCG.R2 | 0.430 | 4.055 | < 0.001 | 0.002 |
| 36 | PreCG.R2–PreCG.L | 0.256 | 4.015 | < 0.001 | 0.003 |
| 37 | IPS.L1–IPL.L2 | 0.396 | 3.994 | < 0.001 | 0.003 |
| 38 | PCun.R1–IC.R | 0.332 | 3.987 | < 0.001 | 0.003 |
| 39 | PCun.R2–IPS.R | 0.388 | 3.919 | < 0.001 | 0.003 |
| 40 | Vermis7–SPL.L | 0.302 | 3.753 | 0.001 | 0.005 |
| 41 | SPL.L–IPL.L2 | 0.556 | 3.740 | 0.001 | 0.005 |
| 42 | MOG.L–SPL.L | 0.425 | 3.739 | 0.001 | 0.005 |
| 43 | SPL.L–PreCG.L | 0.285 | 3.733 | 0.001 | 0.005 |
| 44 | PCun.R1–PreCG.R2 | 0.393 | 3.733 | 0.001 | 0.005 |
| 45 | IPS.R–IPL.L2 | 0.427 | 3.696 | 0.001 | 0.005 |
| 46 | MOG.L–IPS.R | 0.367 | 3.695 | 0.001 | 0.005 |
| 47 | MOG.L–PCun.L | 0.468 | 3.679 | 0.001 | 0.006 |
| 48 | PreCG.R2–SMA.L | 0.494 | 3.611 | 0.001 | 0.006 |
| 49 | PCun.R1–IPL.L1 | 0.408 | 3.600 | 0.001 | 0.007 |
| 50 | IPS.R–IPL.L1 | 0.400 | 3.565 | 0.001 | 0.007 |
| 51 | PCun.R2–PreCG.R1 | 0.338 | 3.559 | 0.001 | 0.007 |
| 52 | CereVI.L–IPS.R | 0.270 | 3.507 | 0.001 | 0.008 |
| 53 | IC.L–PreCG.L | 0.185 | 3.444 | 0.001 | 0.009 |
| 54 | PCun.L–PreCG.R1 | 0.351 | 3.418 | 0.002 | 0.010 |
| 55 | PCun.R1–IPL.L2 | 0.376 | 3.414 | 0.002 | 0.010 |
| 56 | IPS.R–PreCG.R2 | 0.356 | 3.374 | 0.002 | 0.011 |
| 57 | PCun.R1–SMA.L | 0.407 | 3.343 | 0.002 | 0.012 |
| 58 | PCun.R1–IPS.L2 | 0.280 | 3.299 | 0.002 | 0.013 |
| 59 | Vermis7–IPS.R | 0.277 | 3.268 | 0.002 | 0.014 |
| 60 | PCun.L–IC.R | 0.279 | 3.246 | 0.002 | 0.014 |
| 61 | IPS.L1–PreCG.L | 0.217 | 3.190 | 0.003 | 0.016 |
| 62 | PCun.R2–SPL.L | 0.307 | 3.172 | 0.003 | 0.017 |
| 63 | CereVI.L–PreCG.R2 | 0.244 | 3.167 | 0.003 | 0.017 |
| 64 | SMA.L–PreCG.R1 | 0.263 | 3.166 | 0.003 | 0.017 |
| 65 | Cun.L–PCun.R2 | 0.234 | 3.081 | 0.004 | 0.020 |
| 66 | IPL.L1–PreCG.L | 0.229 | 3.071 | 0.004 | 0.021 |
| 67 | CereVII.L–PCun.L | 0.267 | 3.032 | 0.004 | 0.023 |
| 68 | PCun.R2–IC.L | 0.242 | 3.003 | 0.005 | 0.024 |
| 69 | PCun.R2–IPL.L1 | 0.284 | 2.907 | 0.006 | 0.029 |
| 70 | IPS.R–IC.L | 0.237 | 2.880 | 0.006 | 0.031 |
| 71 | Cun.L–PCun.R1 | 0.253 | 2.869 | 0.007 | 0.031 |
| 72 | PCun.R2–PreCG.L | 0.199 | 2.843 | 0.007 | 0.033 |
| 73 | PCun.L–SMA.L | 0.291 | 2.826 | 0.007 | 0.034 |
| 74 | Cun.L–SPL.L | 0.272 | 2.817 | 0.008 | 0.035 |
| 75 | PCun.R1–MCC.R | 0.190 | 2.812 | 0.008 | 0.035 |
| 76 | PreCG.R1–PreCG.L | 0.208 | 2.795 | 0.008 | 0.036 |
| 77 | Thal.L–SMA.L | 0.193 | 2.777 | 0.008 | 0.037 |
| 78 | Put.L1–PreCG.L | 0.170 | 2.767 | 0.009 | 0.038 |
| 79 | IPL.L2–IPL.L1 | 0.326 | 2.762 | 0.009 | 0.038 |
| 80 | Vermis6–IPL.L2 | 0.277 | 2.760 | 0.009 | 0.038 |
| 81 | Vermis7–PreCG.R2 | 0.200 | 2.755 | 0.009 | 0.039 |
| 82 | PCun.L–IC.L | 0.168 | 2.725 | 0.010 | 0.040 |
| 83 | IPL.L1–IC.L | 0.163 | 2.706 | 0.010 | 0.042 |
| 84 | CereVIII.L–PCun.L | 0.248 | 2.681 | 0.011 | 0.043 |
| 85 | PCun.R1–PreCG.L | 0.268 | 2.679 | 0.011 | 0.044 |
| 86 | IC.R–PreCG.L | 0.206 | 2.665 | 0.011 | 0.044 |
| 87 | Vermis9–IC.L | 0.171 | 2.654 | 0.012 | 0.045 |
| 88 | IC.R–PreCG.R1 | 0.215 | 2.652 | 0.012 | 0.045 |
| 89 | IPS.L1–SMA.L | 0.266 | 2.640 | 0.012 | 0.046 |
| 90 | Put.L2–IPS.R | 0.233 | 2.636 | 0.012 | 0.046 |
| 91 | Cun.L–PCun.L | 0.271 | 2.624 | 0.012 | 0.047 |
| 92 | SMA.L–PreCG.L | 0.186 | 2.618 | 0.013 | 0.048 |
| 93 | MOG.L–PreCG.R2 | 0.280 | 2.616 | 0.013 | 0.048 |
| 94 | CereVI.L–PCun.L | 0.253 | 2.599 | 0.013 | 0.049 |

*Note.* All 94 significant ROI-to-ROI connections, ranked by *t*-value (descending). The connection highlighted in bold (MCC.R–HPC.R) represents the primary connection of interest discussed in the main text.

**Supplementary Table 4.** Significant functional connectivity within the map maintenance network (FDR-corrected *p* < 0.05).

| No. | Regions | Connectivity Strength | $t(38)$ | $p$ | $p\_FDR$ |
| --- | --- | --- | --- | --- | --- |
| 1 | MTG.L2–MTG.L1 | 0.391 | 4.887 | < 0.001 | 0.005 |
| 2 | AG.L–ITG.R | 0.431 | 3.962 | < 0.001 | 0.036 |
| 3 | AG.L–MTG.L1 | 0.356 | 3.778 | 0.001 | 0.044 |

**Supplementary Table 5.** Model RDM fits for learning- and updating-related neural ΔRDMs.

| ROI | Phase | Model RDM | Mean Fisher-z fit | $t(42)$ | $p$ | $p\_FDR$ | Cohen's $d$ |
| --- | --- | --- | --- | --- | --- | --- | --- |
| HPC.R | learning | Original | 0.068 | 3.264 | 0.002 | 0.004 | 0.498 |
|  | updating | Original | 0.054 | 3.072 | 0.004 | 0.007 | 0.469 |
|  | updating | Updated | -0.052 | -3.095 | 0.003 | 0.007 | -0.472 |
| HPC.L | learning | Original | 0.018 | 0.820 | 0.417 | 0.417 | 0.125 |
|  | updating | Original | -0.006 | -0.233 | 0.817 | 0.817 | -0.036 |
|  | updating | Updated | -0.027 | -1.323 | 0.193 | 0.193 | -0.202 |

*Note.* Model fits were computed as Spearman correlations between model RDMs and neural ΔRDMs ( ΔRDM<sub>learn</sub>, calculated as PVT post-learn minus PVT pre-learn; ΔRDM<sub>update</sub>, calculated as PVT post-update minus PVT pre-learn) and Fisher $z$ -transformed prior to group-level statistics. FDR correction was applied across the two ROIs within each phase/model combination. Positive fits indicate representational differentiation, whereas negative fits indicate representational integration (see Methods).

**Supplementary Table 6.**
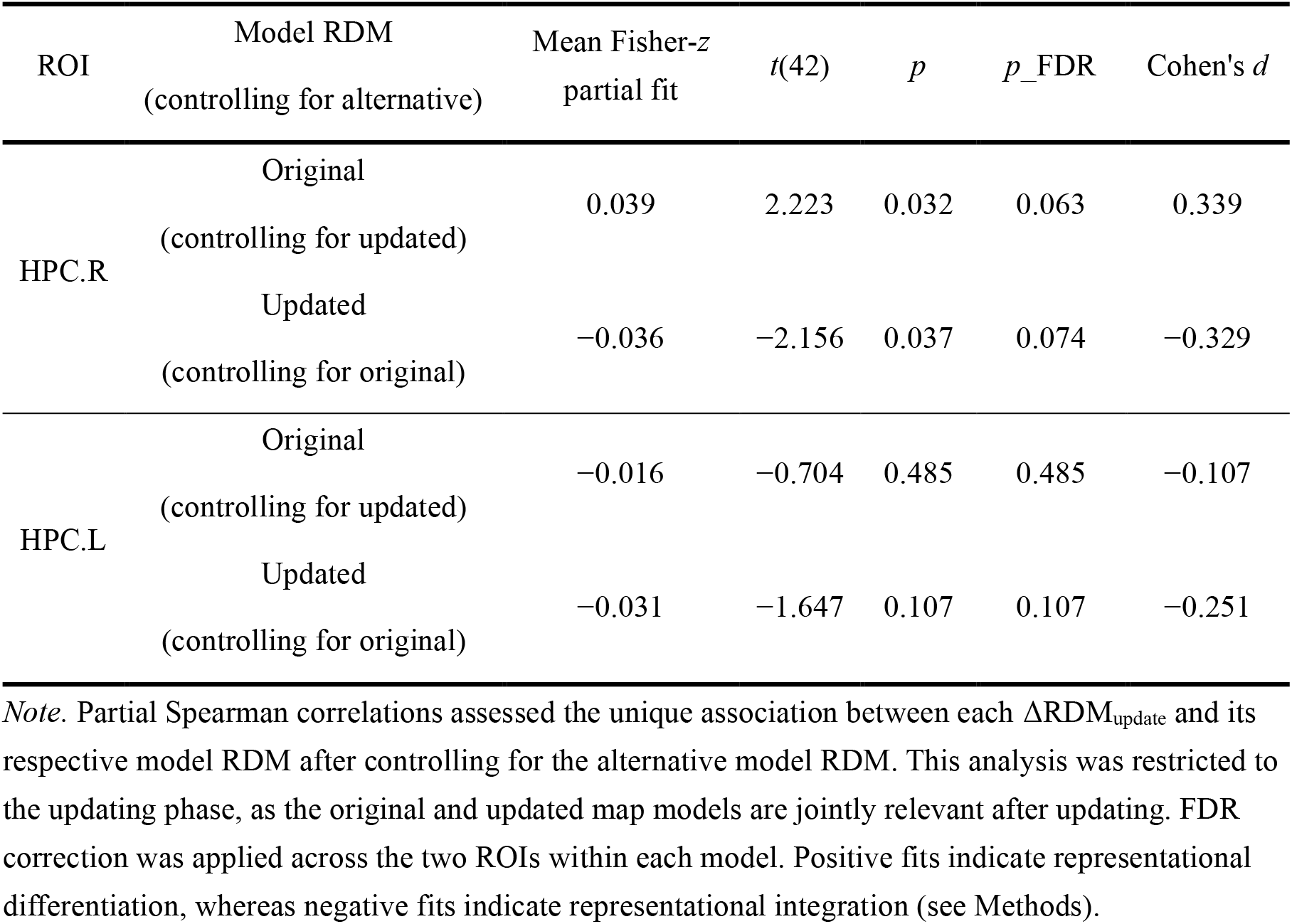
Partial model RDM fits for updating-related neural ΔRDMs.

**Supplementary Table 7.** Association between hippocampal–MCC functional connectivity and hippocampal representational change (ΔW−B) for the updated and original map structures, controlling for mean framewise displacement (mean FD).

| Model | $b$ | 95% CI | SE | $t(35)$ | $p$ | $p\_FDR$ | std $\beta$ | Partial $r$ |
| --- | --- | --- | --- | --- | --- | --- | --- | --- |
| Updated map | -0.008 | [-0.014, -0.002] | 0.003 | -2.721 | 0.010 | 0.020 | -0.414 | -0.418 |
| Original map | 0.003 | [-0.006, 0.011] | 0.004 | 0.669 | 0.508 | 0.508 | 0.113 | 0.112 |

*Note.* $b$ , unstandardized regression coefficient; CI, confidence interval; SE, standard error; std $\beta$ , standardized regression coefficient. Degrees of freedom ( $df = 35$ ) reflect $n = 38$ participants with valid PVT and spatial map updating data, minus 3 estimated parameters (intercept, connectivity, mean FD). Partial $r$ values were computed via bootstrap resampling (5,000 iterations) and used to test the difference between the updated- and original-map associations ( $\Delta r = -0.530$ , 95% CI [-0.912, -0.100], $p = 0.013$ ).

## References

1. Tolman, E. C. Cognitive maps in rats and men. Psychological Review 55, 189–208 (1948).

2. Epstein, R. A., Patai, E. Z., Julian, J. B. & Spiers, H. J. The cognitive map in humans: spatial navigation and beyond. Nat Neurosci 20, 1504–1513 (2017).

3. Peer, M., Brunec, I. K., Newcombe, N. S. & Epstein, R. A. Structuring Knowledge with Cognitive Maps and Cognitive Graphs. Trends in Cognitive Sciences 25, 37–54 (2021).

4. Johnson, A. & Redish, A. D. Neural Ensembles in CA3 Transiently Encode Paths Forward of the Animal at a Decision Point. J. Neurosci. 27, 12176–12189 (2007).

5. Karlsson, M. P. & Frank, L. M. Awake replay of remote experiences in the hippocampus. Nat Neurosci 12, 913–918 (2009).

6. O’Keefe, J. & Nadel, L. Précis of O’Keefe & Nadel’s The hippocampus as a cognitive map. Behav Brain Sci 2, 487–494 (1979).

7. Brown, T. I. et al. Prospective representation of navigational goals in the human hippocampus. Science 352, 1323–1326 (2016).

8. Deuker, L., Bellmund, J. L., Navarro Schröder, T. & Doeller, C. F. An event map of memory space in the hippocampus. eLife 5, e16534 (2016).

9. Behrens, T. E. J. et al. What Is a Cognitive Map? Organizing Knowledge for Flexible Behavior. Neuron 100, 490–509 (2018).

10. Eichenbaum, H. Hippocampus. Neuron 44, 109–120 (2004).

11. Garvert, M. M., Dolan, R. J. & Behrens, T. E. A map of abstract relational knowledge in the human hippocampal–entorhinal cortex. eLife 6, e17086 (2017).

12. O’Keefe, J. & Dostrovsky, J. The hippocampus as a spatial map. Preliminary evidence from unit activity in the freely-moving rat. Brain Research 34, 171–175 (1971).

13. Yang, C. & Naya, Y. Sequential involvements of the perirhinal cortex and hippocampus in the recall of item-location associative memory in macaques. PLoS Biol 21, e3002145 (2023).

14. Bakker, A., Kirwan, C. B., Miller, M. & Stark, C. E. L. Pattern Separation in the Human Hippocampal CA3 and Dentate Gyrus. Science 319, 1640–1642 (2008).

15. Chanales, A. J. H., Oza, A., Favila, S. E. & Kuhl, B. A. Overlap among Spatial Memories Triggers Repulsion of Hippocampal Representations. Current Biology 27, 2307–2317.e5 (2017).

16. Favila, S. E., Chanales, A. J. H. & Kuhl, B. A. Experience-dependent hippocampal pattern differentiation prevents interference during subsequent learning. Nat Commun 7, 11066 (2016).

17. Schlichting, M. L., Mumford, J. A. & Preston, A. R. Learning-related representational changes reveal dissociable integration and separation signatures in the hippocampus and prefrontal cortex. Nat Commun 6, 8151 (2015).

18. Tompary, A. & Davachi, L. Consolidation Promotes the Emergence of Representational Overlap in the Hippocampus and Medial Prefrontal Cortex. Neuron 96, 228–241.e5 (2017).

19. Fernandez, C., Jiang, J., Wang, S.-F., Choi, H. L. & Wagner, A. D. Representational integration and differentiation in the human hippocampus following goal-directed navigation. eLife 12, e80281 (2023).

20. Levy, S. J. & Hasselmo, M. E. Hippocampal remapping induced by new behavior is mediated by spatial context. Preprint at 10.1101/2023.02.20.529330 (2023).

21. Hasselmo, M. E. & Wyble, B. P. Free recall and recognition in a network model of the hippocampus: simulating effects of scopolamine on human memory function. Behavioural Brain Research 89, 1–34 (1997).

22. McClelland, J. L., McNaughton, B. L. & O’Reilly, R. C. Why there are complementary learning systems in the hippocampus and neocortex: insights from the successes and failures of connectionist models of learning and memory. Psychological review 102, 419 (1995).

23. McCloskey, M. & Cohen, N. J. Catastrophic Interference in Connectionist Networks: The Sequential Learning Problem. in Psychology of Learning and Motivation vol. 24 109–165 (Elsevier, 1989).

24. Grossberg, S. How does a brain build a cognitive code? Psychological Review 87, 1–51 (1980).

25. Wang, F. & Bicanski, A. Dynamic Updating of Cognitive Maps via Traces of Experience in the Subiculum. Hippocampus 36, e70078 (2026).

26. Clark, B. J., Simmons, C. M., Berkowitz, L. E. & Wilber, A. A. The retrosplenial-parietal network and reference frame coordination for spatial navigation. Behavioral Neuroscience 132, 416–429 (2018).

27. Zheng (征亦诚), Y., et al. A Hippocampal–Parietal Network for Reference Frame Coordination. J. Neurosci. 45, e1782242025 (2025).

28. Schlichting, M. L. & Preston, A. R. Hippocampal–medial prefrontal circuit supports memory updating during learning and post-encoding rest. Neurobiology of Learning and Memory 134, 91–106 (2016).

29. Domic-Siede, M., Irani, M., Valdés, J., Perrone-Bertolotti, M. & Ossandón, T. Theta activity from frontopolar cortex, mid-cingulate cortex and anterior cingulate cortex shows different roles in cognitive planning performance. NeuroImage 226, 117557 (2021).

30. Huster, R. J., Enriquez-Geppert, S., Pantev, C. & Bruchmann, M. Variations in midcingulate morphology are related to ERP indices of cognitive control. Brain Struct Funct 219, 49–60 (2014).

31. Kwapis, J. L. et al. Aging mice show impaired memory updating in the novel OUL updating paradigm. Neuropsychopharmacol. 45, 337–346 (2020).

32. Wright, D. S., Bodinayake, K. K. & Kwapis, J. L. Investigating Memory Updating in Mice Using the Objects in Updated Locations Task. CP Neuroscience 91, e87 (2020).

33. Fukushima, T., Hasegawa, I. & Miyashita, Y. Prefrontal Neuronal Activity Encodes Spatial Target Representations Sequentially Updated After Nonspatial Target-Shift Cues. Journal of Neurophysiology 91, 1367–1380 (2004).

34. Ottink, L., De Haas, N. & Doeller, C. F. Integration of Euclidean and path distances in hippocampal maps. Sci Rep 15, 7104 (2025).

35. Hulbert, J. C. & Norman, K. A. Neural Differentiation Tracks Improved Recall of Competing Memories Following Interleaved Study and Retrieval Practice. Cereb. Cortex 25, 3994–4008 (2015).

36. LaRocque, K. F. et al. Global Similarity and Pattern Separation in the Human Medial Temporal Lobe Predict Subsequent Memory. J. Neurosci. 33, 5466–5474 (2013).

37. Yassa, M. A. & Stark, C. E. L. Pattern separation in the hippocampus. Trends in Neurosciences 34, 515–525 (2011).

38. Schlichting, M. L. & Preston, A. R. Memory integration: neural mechanisms and implications for behavior. Current Opinion in Behavioral Sciences 1, 1–8 (2015).

39. Molitor, R. J., Sherrill, K. R., Morton, N. W., Miller, A. A. & Preston, A. R. Memory Reactivation during Learning Simultaneously Promotes Dentate Gyrus/CA2,3 Pattern Differentiation and CA1 Memory Integration. J. Neurosci. 41, 726–738 (2021).

40. Mau, W., Hasselmo, M. E. & Cai, D. J. The brain in motion: How ensemble fluidity drives memory-updating and flexibility. eLife 9, e63550 (2020).

41. Iglói, K., Doeller, C. F., Berthoz, A., Rondi-Reig, L. & Burgess, N. Lateralized human hippocampal activity predicts navigation based on sequence or place memory. Proc. Natl. Acad. Sci. U.S.A. 107, 14466–14471 (2010).

42. Piekema, C., Kessels, R. P. C., Mars, R. B., Petersson, K. M. & Fernández, G. The right hippocampus participates in short-term memory maintenance of object–location associations. NeuroImage 33, 374–382 (2006).

43. Jerde, T. A., Merriam, E. P., Riggall, A. C., Hedges, J. H. & Curtis, C. E. Prioritized Maps of Space in Human Frontoparietal Cortex. J. Neurosci. 32, 17382–17390 (2012).

44. Vossel, S., Geng, J. J. & Fink, G. R. Dorsal and Ventral Attention Systems: Distinct Neural Circuits but Collaborative Roles. Neuroscientist 20, 150–159 (2014).

45. Cavanna, A. E. & Trimble, M. R. The precuneus: a review of its functional anatomy and behavioural correlates. Brain 129, 564–583 (2006).

46. Dordevic, M., Hoelzer, S., Russo, A., García Alanis, J. C. & Müller, N. G. The Role of the Precuneus in Human Spatial Updating in a Real Environment Setting—A cTBS Study. Life 12, 1239 (2022).

47. Stoodley, C. J., Valera, E. M. & Schmahmann, J. D. Functional topography of the cerebellum for motor and cognitive tasks: An fMRI study. NeuroImage 59, 1560–1570 (2012).

48. Wandell, B. A., Dumoulin, S. O. & Brewer, A. A. Visual Field Maps in Human Cortex. Neuron 56, 366–383 (2007).

49. Kravitz, D. J., Saleem, K. S., Baker, C. I., Ungerleider, L. G. & Mishkin, M. The ventral visual pathway: an expanded neural framework for the processing of object quality. Trends in Cognitive Sciences 17, 26–49 (2013).

50. Botvinick, M. M., Cohen, J. D. & Carter, C. S. Conflict monitoring and anterior cingulate cortex: an update. Trends in Cognitive Sciences 8, 539–546 (2004).

51. Parvaz, M. A. et al. Multimodal evidence of regional midcingulate gray matter volume underlying conflict monitoring. NeuroImage: Clinical 5, 10–18 (2014).

52. Vogt, B. A. Cingulate cortex in the three limbic subsystems. in Handbook of Clinical Neurology vol. 166 39–51 (Elsevier, 2019).

53. Kriegeskorte, N. Representational similarity analysis – connecting the branches of systems neuroscience. Front. Sys. Neurosci. https://doi.org/10.3389/neuro.06.004.2008 (2008) doi:10.3389/neuro.06.004.2008.

54. Cocchi, L., Zalesky, A., Fornito, A. & Mattingley, J. B. Dynamic cooperation and competition between brain systems during cognitive control. Trends in Cognitive Sciences 17, 493–501 (2013).

55. Shine, J. M. et al. The Dynamics of Functional Brain Networks: Integrated Network States during Cognitive Task Performance. Neuron 92, 544–554 (2016).

56. Tse, D. et al. Schema-Dependent Gene Activation and Memory Encoding in Neocortex. Science 333, 891–895 (2011).

57. Van Kesteren, M. T. R., Fernández, G., Norris, D. G. & Hermans, E. J. Persistent schema-dependent hippocampal-neocortical connectivity during memory encoding and postencoding rest in humans. Proc. Natl. Acad. Sci. U.S.A. 107, 7550–7555 (2010).

58. Van Kesteren, M. T. R., Ruiter, D. J., Fernández, G. & Henson, R. N. How schema and novelty augment memory formation. Trends in Neurosciences 35, 211–219 (2012).

59. Van Kesteren, M. T. R. et al. Differential roles for medial prefrontal and medial temporal cortices in schema-dependent encoding: From congruent to incongruent. Neuropsychologia 51, 2352–2359 (2013).

60. Hsu, N. S., Schlichting, M. L. & Thompson-Schill, S. L. Feature Diagnosticity Affects Representations of Novel and Familiar Objects. Journal of Cognitive Neuroscience 26, 2735–2749 (2014).

61. Willenbockel, V. et al. Controlling low-level image properties: The SHINE toolbox. Behavior Research Methods 42, 671–684 (2010).

62. Esteban, O. et al. fMRIPrep: a robust preprocessing pipeline for functional MRI. Nat Methods 16, 111–116 (2019).

63. Gorgolewski, K. et al. Nipype: A Flexible, Lightweight and Extensible Neuroimaging Data Processing Framework in Python. Front. Neuroinform. 5, (2011).

64. Tustison, N. J. et al. N4ITK: Improved N3 Bias Correction. IEEE Trans. Med. Imaging 29, 1310–1320 (2010).

65. Avants, B., Epstein, C., Grossman, M. & Gee, J. Symmetric diffeomorphic image registration with cross-correlation: Evaluating automated labeling of elderly and neurodegenerative brain. Medical Image Analysis 12, 26–41 (2008).

66. Zhang, Y., Brady, M. & Smith, S. Segmentation of brain MR images through a hidden Markov random field model and the expectation-maximization algorithm. IEEE Trans. Med. Imaging 20, 45–57 (2001).

67. Dale, A. M., Fischl, B. & Sereno, M. I. Cortical Surface-Based Analysis. NeuroImage 9, 179–194 (1999).

68. Ciric, R. et al. TemplateFlow: FAIR-sharing of multi-scale, multi-species brain models. Nat Methods 19, 1568–1571 (2022).

69. Pruim, R. H. R. et al. ICA-AROMA: A robust ICA-based strategy for removing motion artifacts from fMRI data. NeuroImage 112, 267–277 (2015).

70. Lohmann, G. et al. LISA improves statistical analysis for fMRI. Nat Commun 9, 4014 (2018).

71. Ness, H. T. et al. Reduced Hippocampal-Striatal Interactions during Formation of Durable Episodic Memories in Aging. Cerebral Cortex bhab331 (2021) doi:10.1093/cercor/bhab331.

72. Schiller, D. et al. Memory and Space: Towards an Understanding of the Cognitive Map. J. Neurosci. 35, 13904–13911 (2015).

73. Walther, A. et al. Reliability of dissimilarity measures for multi-voxel pattern analysis. NeuroImage 137, 188–200 (2016).

74. McLaren, D. G., Ries, M. L., Xu, G. & Johnson, S. C. A generalized form of context-dependent psychophysiological interactions (gPPI): A comparison to standard approaches. NeuroImage 61, 1277–1286 (2012).

75. Nieto-Castanon, A. Handbook of Functional Connectivity Magnetic Resonance Imaging Methods in CONN. (Hilbert Press, 2020). doi:10.56441/hilbertpress.2207.6598.

76. Behzadi, Y., Restom, K., Liau, J. & Liu, T. T. A component based noise correction method (CompCor) for BOLD and perfusion based fMRI. NeuroImage 37, 90–101 (2007).

77. Power, J. D. et al. Methods to detect, characterize, and remove motion artifact in resting state fMRI. NeuroImage 84, 320–341 (2014).

78. Alakörkkö, T., Saarimäki, H., Glerean, E., Saramäki, J. & Korhonen, O. Effects of spatial smoothing on functional brain networks. Eur J of Neuroscience 46, 2471–2480 (2017).

79. Strimmer, K. A unified approach to false discovery rate estimation. BMC Bioinformatics 9, 303 (2008).

80. Strimmer, K. fdrtool: a versatile R package for estimating local and tail area-based false discovery rates. Bioinformatics 24, 1461–1462 (2008).

